# The differentiation and functional landscape of tumor-resident CD8^+^ T cells shapes immunity and clinical outcomes in clear cell renal cell carcinoma

**DOI:** 10.64898/2026.03.12.711359

**Authors:** Sofía Hidalgo, Andrés Hernández-Oliveras, Jimena Tosello-Boari, Javiera Reyes-Alvarez, Sergio Hernández-Galaz, Catalina Bustamante, Ernesto López, Yoann Missolo-Koussou, Wilfrid Richer, Vincenzo Benedetti, Farides Saavedra, Marco Fraga, Eduardo Roa, Diego Figueroa, Christine Sedlik, Jordan Denizeau, Luis Alarcón, Felipe Gálvez-Cancino, Daniela Sauma, Thierry Lebret, Camelia Radulescu, Yves Allory, Vincenzo Borgna, Eliane Piaggio, Alvaro Lladser

## Abstract

Clear cell renal cell carcinoma (ccRCC) is a common and lethal kidney cancer in which CD8^+^ T cell–targeted immunotherapies are standard of care. However, tumor CD8^+^ T cell infiltration correlates with divergent clinical outcomes, reflecting functional heterogeneity. We comprehensively characterized tumor-specific memory CD8^+^ T cells in ccRCC using phenotypic, transcriptional and functional analyses. Three major subsets were identified based on tissue-residency markers: circulating (Tcirc), tissue-resident (Trm) and CD69^+^CD103^−^ Trm-like cells, all recognizing autologous RCC cells in an HLA class I–dependent manner. Tcirc cells exhibited greater stemness and cytotoxic potential, whereas Trm populations displayed tissue-residency programs, enhanced tumor reactivity and exhaustion features. Single-cell transcriptomics analyses uncovered substantial heterogeneity within Trm cells, including progenitor, activated, transitional, type-17, interferon-responsive, regulatory and terminally exhausted states. Trajectory and TCR analyses indicated progressive differentiation from central-memory Tcirc cells toward tissue residency while losing circulation, cytotoxicity and stemness potentials and then progress toward terminal exhaustion. Enrichment of progenitor Trm cells associated with improved survival and favorable immunotherapy response, whereas regulatory or exhausted Trm cells correlated with poor outcomes. These findings define the differentiation and functional landscape of tumor-specific CD8^+^ T cells that shapes immunity and clinical outcomes in ccRCC.

## Main

Clear cell renal cell carcinoma (ccRCC) accounts for approximately 83-88% of kidney cancer cases and the majority of kidney-cancer related deaths^1^. The 5-year survival rates for localized, regional and distant ccRCC are 93%, 70% and 13%, respectively^2^. Moreover, nearly 50% of patients with localized tumors develop metastases or experience recurrence after nephrectomy^3^. ccRCC tumors are highly infiltrated by T cells, comprising on average nearly half of the immune population within the tumor microenvironment^4^. In contrast to most malignancies, CD8^+^ T cell infiltration in primary and metastatic ccRCC correlates with poor survival, a paradox likely explained by the dysfunctional state of tumor-infiltrating CD8^+^ T cells^4–7^.

Immunotherapy using immune checkpoint inhibitors (ICI), which reinvigorates T cell function by blocking inhibitory receptors, has revolutionized cancer treatment. Agents such as pembrolizumab and nivolumab (anti-PD-1), avelumab (anti-PD-L1), and ipilimumab (anti-CTLA4), alone or in combination with other therapies, significantly improved clinical outcomes in ccRCC^8–11^. Despite this beneficial effect, a significant fraction of patients fails to respond or develop resistance to ICI. Comprehensive characterization of CD8^+^ T cell populations that drive clinical benefit from immunotherapy remains incomplete.

Recent single-cell genomics studies have begun to uncover the composition and dynamics of tumor-infiltrating T cells that influence response or resistance to ICI^12,13^. Memory CD8^+^ T cells are considered key actors in cancer immunotherapy due to their capacity to exert durable responses compared to short-lived effector CD8^+^ T cells^14^. These cells differentiate into circulating and tissue-resident memory compartments: circulating cells patrol blood, lymphoid and peripheral organs, whereas tissue-resident cells remain localized within tumor sites. Several studies have revealed that together, circulating (Tcirc) and resident memory (Trm) CD8^+^ T cells orchestrate durable immunity across different cancers^14,15^.

Particularly, Trm cells have been linked to favorable immunotherapy responses in various types of cancer, including lung, cervical, colorectal, esophageal, gastric, head and neck, and melanoma^16,17^. In ccRCC, expansion of Trm cells has been associated with favorable responses to ICI, and a Trm cell-derived gene signature predicts better progression-free survival in patients treated with avelumab plus axitinib^18^. In contrast, CD8^+^ T cells exhibiting terminally exhausted phenotypes correlate with cancer progression and poor survival^19,20^. However, direct evidence of tumor reactivity among memory CD8^+^ T cell subsets remains limited, and previous single-cell studies lack sufficient resolution to fully capture their differentiation and functional diversity, limiting our understanding of their clinical relevance.

Here, we comprehensively characterize the phenotypic, functional, and transcriptional diversity of tumor-infiltrating memory CD8^+^ T cells in ccRCC, uncovering differentiation trajectories and their association with clinical outcomes. We identify three tumor-specific memory CD8^+^ T cell populations in ccRCC, Tcirc, Trm, and CD69^+^CD103^−^ Trm-like cells, with distinct differentiation states and tumor reactivity. Tcirc cells retain stemness, while Trm populations show tissue adaptation and progressive exhaustion. Progenitor Trm enrichment predicts improved survival and immunotherapy response, whereas exhausted Trm and total CD8^+^ T cell abundance associate with poor outcomes. These findings clarify the differentiation trajectory of tumor-specific CD8^+^ T cells in ccRCC and its clinical relevance.

## Results

### Tumor-infiltrating Trm cells exhibit high reactivity and effector function towards autologous ccRCC cells

To characterize in depth the phenotypic, functional, and transcriptomic heterogeneity of antigen-specific CD8^+^ T cells, we analyzed memory CD8^+^ T cells from primary tumors from untreated ccRCC patients using spectral flow cytometry, coculture assays, bulk and single-cell transcriptomics coupled with TCR sequencing. (**Fig. 1A**). Based on the expression of tissue residency canonical markers CD69 and CD103^21^, memory CD8^+^ T cells infiltrating human ccRCC tumors were classified in three main populations: CD69^−^CD103^−^ Tcirc cells (mean of frequency: 17.9%), CD69^+^CD103^−^ cells (mean of frequency: 52.2%) and CD69^+^CD103^+^ Trm cells (mean of frequency: 26.5%) (**Fig. 1B**). Both CD69^+^CD103^−^ and Trm subpopulations, were enriched in tumors but not peripheral blood (**Fig. 1B-C**), indicative of their tissue residency. The three populations were also found in non-malignant renal tissue (NMRT) adjacent to RCC tumors, with higher proportion of Tcirc cells (mean of frequency: 40.25%) and reduced frequencies of CD69^+^CD103^−^ Trm-like cells (mean of frequency: 32.29%), as compared to tumors (**Supplementary Fig. 1A-B**).

**Figure 1.**
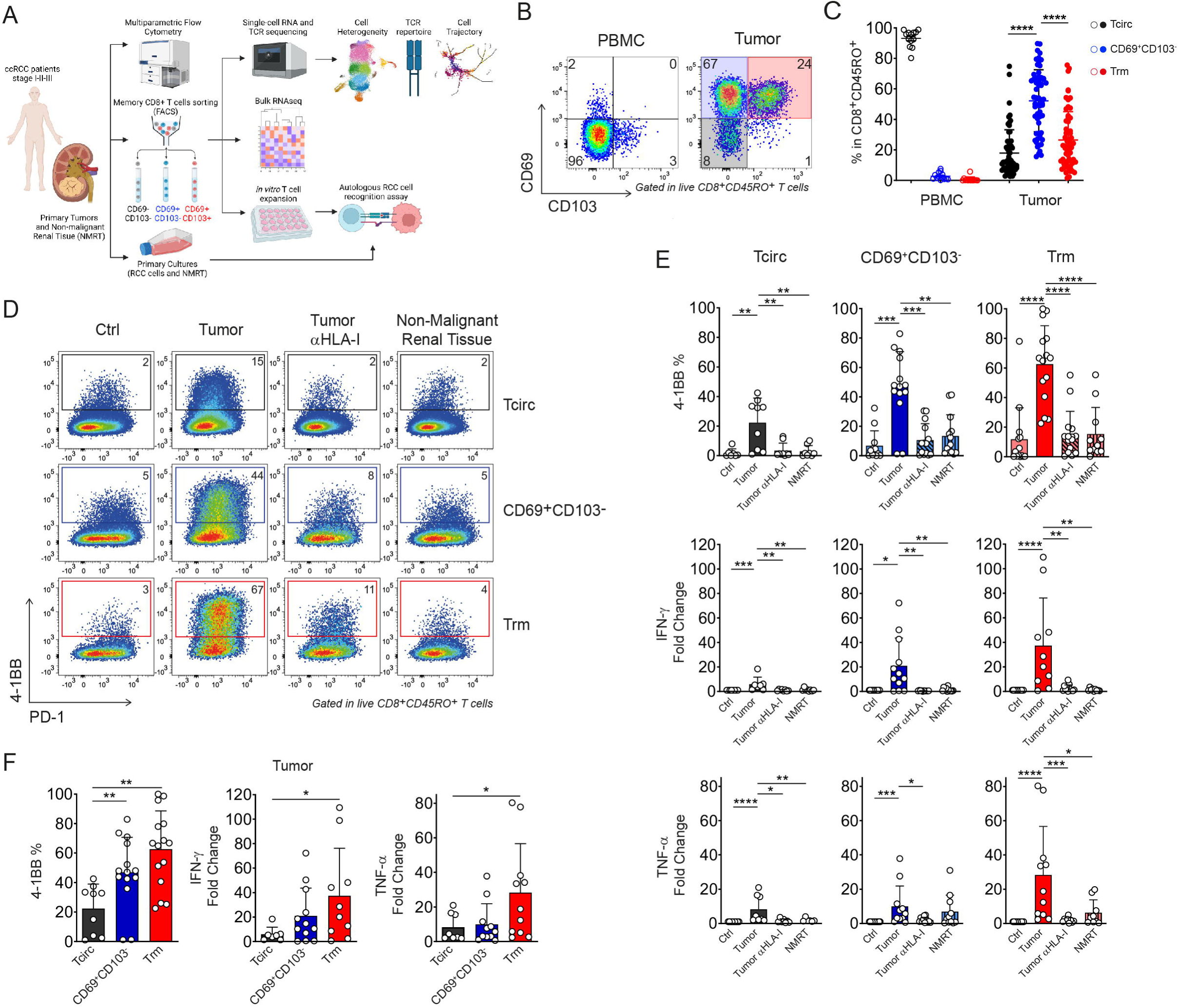
Tumor-infiltrating Trm cells exhibit high reactivity and effector function towards autologous ccRCC cells. **(A)** Samples workflow of the study. **(B)** Representative FACS plots showing CD69 and CD103 expression in CD45RO^+^CD8^+^ TILs from ccRCC samples PBMC isolated from peripheral blood of patients were used as control. **(C)** Frequency quantification of memory CD8^+^ T cells subpopulations. Each point represents a patient. Statistical analysis was performed using Friedman paired one-way ANOVA test. *p < 0.05; **p < 0.01; ***p < 0.001; **** p < 0.0001. **(D-F)** Sorted CD69^−^CD103^−^(Tcirc), CD69^+^CD103^−^ and CD69^+^CD103^+^ (Trm) memory CD8^+^ TIL were expanded in vitro, and tumor reactivity was assessed by coculture with autologous tumor +/- HLA-I blocking antibody, or non-malignant renal tissue (NMRT). **(D)** Representative FACS plot showing 4-1BB upregulation upon ccRCC cell recognition. **(E)** Quantification of 4-1BB upregulation (upper panels), IFN-γ secretion (middle panels) and TNF-α secretion (bottom panels) in each coculture condition for each CD8^+^ TIL subpopulation. **(F)** Comparison of the magnitude of tumor recognition (4-1BB upregulation) and cytokine secretion (IFN-γ and TNF-α) between memory CD8^+^ TIL subpopulation. Each point represents a patient. Statistical analysis was performed using a non-parametrical Mann-Whitney test. *p < 0.05; **p < 0.01; ***p < 0.001; **** p < 0.0001.

Next, we evaluated the ability of the different subsets of memory CD8^+^ T cells to recognize autologous ccRCC cells in coculture experiments. To this end, we isolated Tcirc, CD69^+^CD103^−^Trm-like and Trm subpopulations by cell sorting and expanded them in vitro (**Fig. 1A**). Most of the cells in each subset retained the initial phenotype, with a proportion of Tcirc cells acquiring CD69 expression and a fraction of CD69^+^CD103^−^ Trm-like cells losing CD69 expression, likely reflecting a transient activation state of each subset at the analysis (**Supplementary Fig. 2A-B**). Neither of these two subsets acquire CD103 expression, suggesting that its expression is imprinted in the tissue^22^. A fraction of Trm cells became CD69^+^CD103^−^ but almost none become double negative, indicative of a directional differentiation. In parallel, we generated primary cultures of ccRCC cells and epithelial cells from NMRT, showing a >95% purity assessed by measuring the expression of the epithelial cell marker EpCAM by flow cytometry (**Supplementary Fig. 2C**). The expanded memory CD8^+^ T cell subpopulations were cocultured with autologous ccRCC cells or NMRT and specific recognition was measured by 4-1BB upregulation and the production of the effector molecules IFN-γ and TNF-α^23,24^. The three subpopulations, Tcirc and CD69^+^CD103^−^ Trm-like and Trm cells, upregulated 4-1BB expression and produced IFN-γ and TNF-α in an HLA-I-dependent manner following cocultured with autologous ccRCC cells but not NMRT cells (**Fig. 1D-E**). These results suggest that memory CD8^+^ T cells recognize tumor-specific antigens and not self-antigens expressed in renal tissue. In addition, we observed that CD69^+^CD103^−^ Trm-like and Trm cells exhibited higher tumor reactivity than Tcirc cells (**Fig.1F, left panel**). Interestingly, Trm cells produced significantly higher IFN-γ and TNF-α levels after tumor recognition compared to Tcirc and CD69^+^CD103^−^ Trm-like cells (**Fig. 1F, middle and right panel**). Moreover, only Trm cells produced significant amounts of granzyme B and perforin, indicative of a more functional state (**Supplementary Fig. 2D-E**). These results demonstrate that tumor-infiltrating memory CD8^+^ T cells specifically recognize autologous ccRCC with Trm cells showing higher reactivity and functionality.

### Trm cells constitute a bona fide tissue-resident memory subset related to terminally exhausted CD69^+^CD103^−^ Trm-like cells and distinct from stem-like cytotoxic Tcirc cells

Next, we characterized the three memory CD8^+^ T cell subsets by bulk RNA-seq. Unsupervised principal component analysis (PCA) segregated samples by subset identity (**Supplementary Fig. 3A-B**). PC1, which explained 57% of the variance, separated Tcirc from the CD69^+^CD103^−^ Trm-like and Trm subsets (**Supplementary Fig. 3A**), while patient identity showed a heterogeneous distribution (**Supplementary Fig. 3B**). Across the three populations we identified 5,166 differentially expressed genes (DEGs) **(Fig. 2A**; FDR < 0.05), comprising both unique and shared programs. Trm and CD69^+^CD103^−^ Trm-like cells shared a higher number of DEGs compared with Tcirc cells, consistently with their closely related phenotypes. Tcirc cells exhibited higher expression of recirculation genes (*KLF2*, *S1PR1*, *S1PR5*, CD62L [*SELL*]), together with cytotoxic (perforin [*PRF1*], granulysin [*GNLY*], *CX3CR1*, *FCGR3A*) and stemness-associated (*TCF7*, *IL7R*) transcripts (**Fig. 2B-C and Supplementary Fig. 3C**)^25,26^. In contrast, Trm cells displayed a canonical residency transcription program including CD103 *ITGAE*), Hobit (*ZNF683*), *RBPJ*, *CD101*, and *XCL2*, consistent with tissue establishment^27^. CD69^+^CD103^−^ Trm-like cells expressed the highest levels of inhibitory receptors, including CD39 (*ENTPD1*) and TIM-3 (*HAVCR2*), as well as the transcription factor *TOX* (**Fig. 2B-C and Supplementary Fig. 3C**), indicative of a more terminally differentiated state^28^. Additionally, Trm and CD69^+^CD103^−^Trm-like cells shared the expression of a set of residency-associated genes, such as *ITGA1* (CD49a)*, CXCL13, CXCR6,* and *XCL1* (**Fig. 2B-C and Supplementary Fig. 3C**), indicating an overlapping transcriptional Trm profile^29^. Correlation analysis supported these patterns. Trm-defining markers (*ITGAE*, *ITGA1, XCL1, CD69, TGFBR2, CXCR6*) positively correlated with exhaustion/effector-associated transcripts (*TOX*, *LAYN*, *CXCL13*, *GZMH*, *GZMA, TNFRSF4, TNFRSF18*) and inversely with circulation/cytotoxic-signature genes (*KLF2*, *SELL*, *S1PR1*, *S1PR5*, *CX3CR1*, *FCGR3A*) (**Supplementary Fig. 3D**). We confirmed this characterization using published gene signatures and analyzed by Gene Set Enrichment Analysis (GSEA) in each memory CD8^+^ T cell subpopulations (**Fig. 2D**). We found that previously published precursor-resident and precursor-exhausted (*Gueguen* Prec-Trm and *Li* Prec-Exh) and canonical Trm cell signatures (*Guo* Trm, *Braun* Trm, *Sun* Trm) were enriched in Trm cells, whereas cytotoxic, effector memory (Tem) and progenitor signatures (*Braun* Cytotox, *Anadon* Stem-Tcirc, and *Savas* Tem) were enriched in Tcirc cells (**Fig. 2D**)^19,30,31^. CD69^+^CD103^−^ Trm-like subset transcriptional profiles were associated with exhausted and terminally exhausted cells signature from RCC tumors (*Braun* Term Exh/PMCH^+^Term Exh) and a Trm signature from breast cancer with high levels of inhibitory receptors (*Savas* Trm), indicating a terminally differentiated state (**Fig. 2D**)^19,32^. These results indicate that Trm cells and CD69^+^CD103^−^ Trm-like cells share a residency transcriptional program, with the latter displaying a more terminal stage of differentiation.

**Figure 2.**
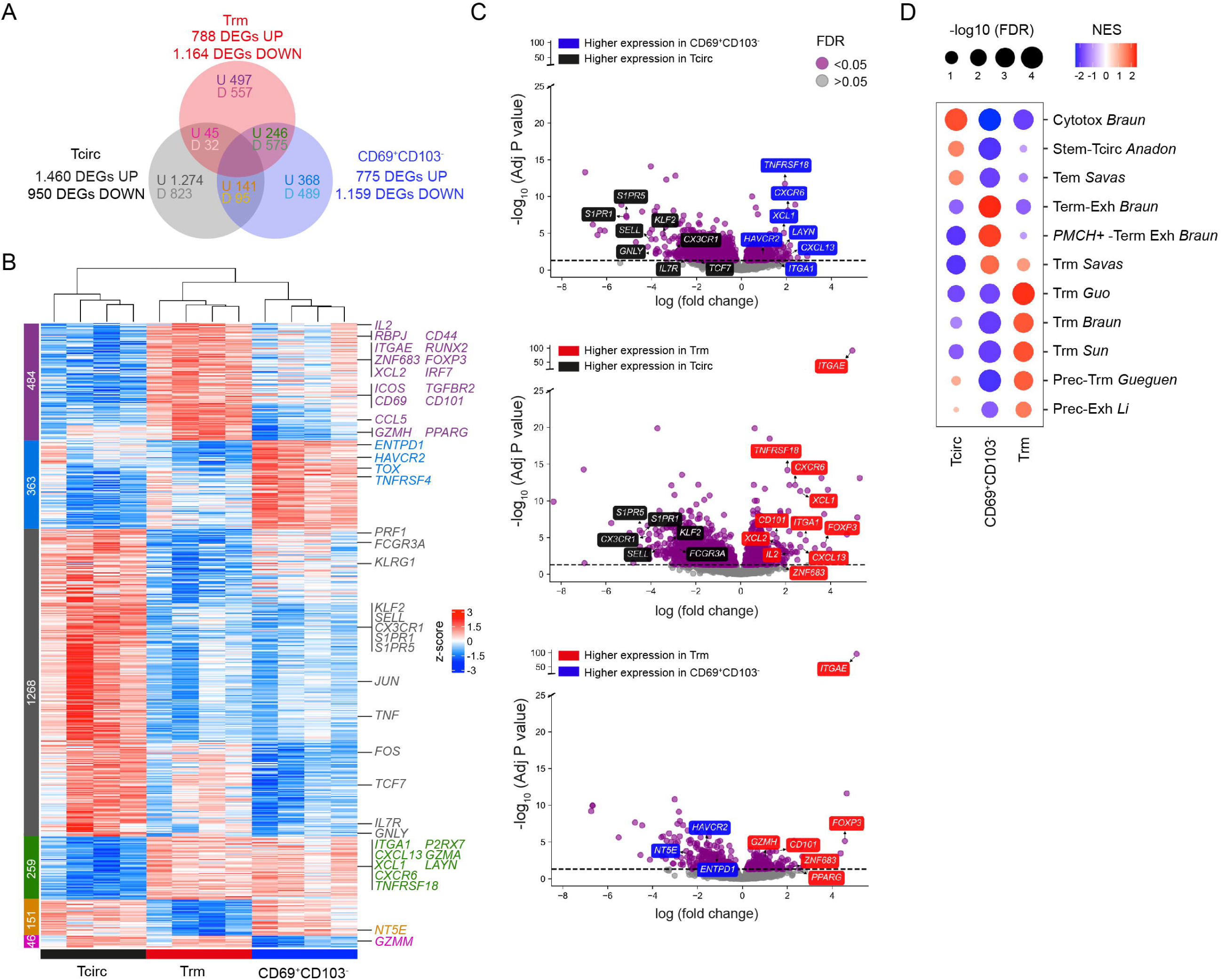
Trm cells constitute a bona fide tissue-resident memory subset related to terminally exhausted CD69^+^CD103^−^ Trm-like cells and distinct from stem-like cytotoxic Tcirc cells. Tcirc, CD69^+^CD103^−^ and Trm cells were isolated from ccRCC tumors and analyzed by bulk RNAseq. **(A)** Venn diagram showing differential expressed genes shared between memory CD8^+^ TILs subpopulations. U= upregulated and D= downregulated. Only genes consistently up- or down-regulated across comparisons are shown. **(B)** Heatmap showing differential expressed genes between Tcirc, CD69^+^CD103- and Trm cells. Key genes are labeled. Genes with positive log2-fold change and adjusted p value < 0.05 were considered as upregulated. Genes with negative log2-fold change and adjusted p value < 0.05 were considered as downregulated. **(C)** Volcano plots showing key markers upregulated in each memory CD8^+^ T cells subpopulation. False discovery rate (FDR) <0.05 was considered significant. **(D)** Gene Set Enrichment Analysis (GSEA) for published gene-expression signatures enriched in Tcirc, CD69^+^CD103^−^ and Trm cells. A signature with positive Normalized Enriched Score (NES) and a -log10-False Discovery Rate (FDR) > 1.3 was considered as positively enriched. A signature with negative NES and a -log10-FDR > 1.3 was considered as negatively enriched.

We validated the transcriptomic profiles of the three populations by flow cytometry. We found that tissue residency-associated integrin CD49a, was highly expressed in Trm and CD69^+^CD103^−^ Trm-like cells, but lowly expressed by Tcirc cells (**Fig. 3A-B**). PD-1 was highly expressed by both CD69^+^CD103^−^ Trm-like and Trm cells and at lower and variable levels by Tcirc cells. In addition, markers associated with tumor antigen reactivity, such as CXCL13 and CD39 were higher at both CD69^+^CD103^−^ and Trm cells (**Fig. 3A-B**). Exhaustion-associated markers CD39, TIM-3 and VCAM-1 were predominantly expressed by CD69^+^CD103^−^ cells but also present in Trm cells. In accordance, the exhaustion-associated transcription factor TOX-1 was highly expressed by these subsets (**Fig. 3C-D**). In contrast, the transcription factor TCF-1 associated with stemness was predominantly present in Tcirc cells, supporting the progenitor status of this subset (**Fig. 3C-D**). Interestingly, in NMRT, all memory CD8^+^ T cell subsets expressed mainly TCF-1 but did not express TOX-1 and accordingly expressed significant lower levels of exhaustion-associated molecules (**Supplementary Fig. 4A-C**), consistent with lack of chronic antigen stimulation. These results suggest that NMRT acts as a reservoir niche of the memory CD8^+^ T cell pool. Finally, we analyzed the expression of cytotoxic molecules and found that even if a high proportion of the three memory CD8^+^ T cell subsets in the tumor expressed granzyme B, only Tcirc cells co-expressed granzyme B and granulysin (**Fig. 3E-F**), indicative of a full cytotoxic potential. These results indicate that memory CD8^+^ T cells infiltrating ccRCC tumors show distinct differentiation subsets, with Tcirc cells exhibiting higher progenitor and cytotoxic potential, bona fide Trm cells displaying a canonical tissue-resident memory program, and CD69^+^CD103^−^ Trm-like showing features of terminal differentiation.

**Figure. 3.**
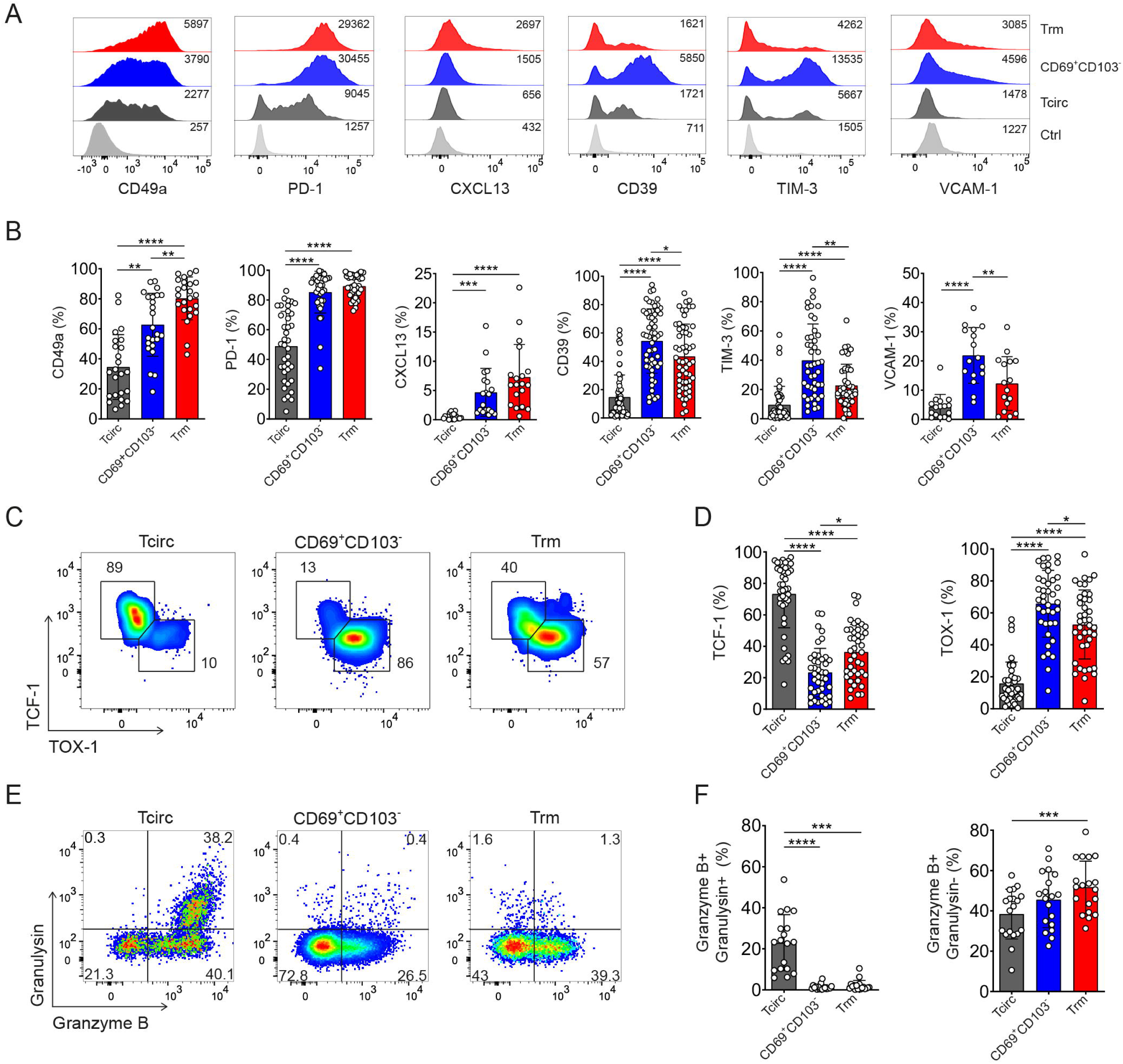
Differential expression of tissue-resident, stemness, cytotoxic and exhaustion associated markers between Tcirc, CD69^+^CD103^−^ Trm-like and Trm cells. **(A-F)** Memory CD8^+^ T cells populations from ccRCC tumors were analyzed by multiparametric flow cytometry and characterized by manual gating. **(A-B)** Histogram plots showing MFI **(A)** and frequency quantification **(B)** for the expression of key markers in memory CD8^+^ T cell subpopulations. **(C)** FACS plots showing TCF1 and TOX expression measured by intranuclear transcription factor staining. **(D)** Frequency of TCF-1 and TOX expression in memory CD8^+^ TIL subpopulations. **(E)** FACS plots showing granzyme B and granulysin intracellular expression **(F)** Frequency quantifications in memory CD8^+^ TIL subpopulations. Each point represents a patient. Statistical analysis was performed using Friedman paired one-way ANOVA test. *p < 0.05; **p < 0.01; ***p < 0.001; **** p < 0.0001.

### High-dimensional analyses reveal heterogenous differentiation states of Trm cells

To gain deeper insights into memory transcriptomic programs, we analyzed tumor-infiltrating memory CD8^+^ T cells by scRNAseq coupled to scTCRseq from three ccRCC patients. After filtering out low-quality cells, we obtained transcriptomic data from 40,721 cells, performed unsupervised graph-based clustering and identified 17 clusters (**Supplementary Fig. 5A, resolution 0.5**). The clusters are displayed in a UMAP using the top 30 principal components (PCs) (**Fig. 4A**). All clusters were present in all patients at varying frequencies (**Supplementary Fig. 5B**) and expressed *CD3E* and *CD8A* but no *CD4* (**Supplementary Fig. 5C**). To annotate clusters, we used: (1) differential expression analysis, (2) gene signatures derived from literature^12,13,19,30–37^, (3) GSEA of scRNAseq cluster signatures in bulk RNA-seq data, and (4) CD69 and CD103 expression analysis using oligonucleotide-barcoded antibodies (antibody-derived tags) (**Fig. 4B-E and Supplementary Fig. 5D**).

**Figure 4.**
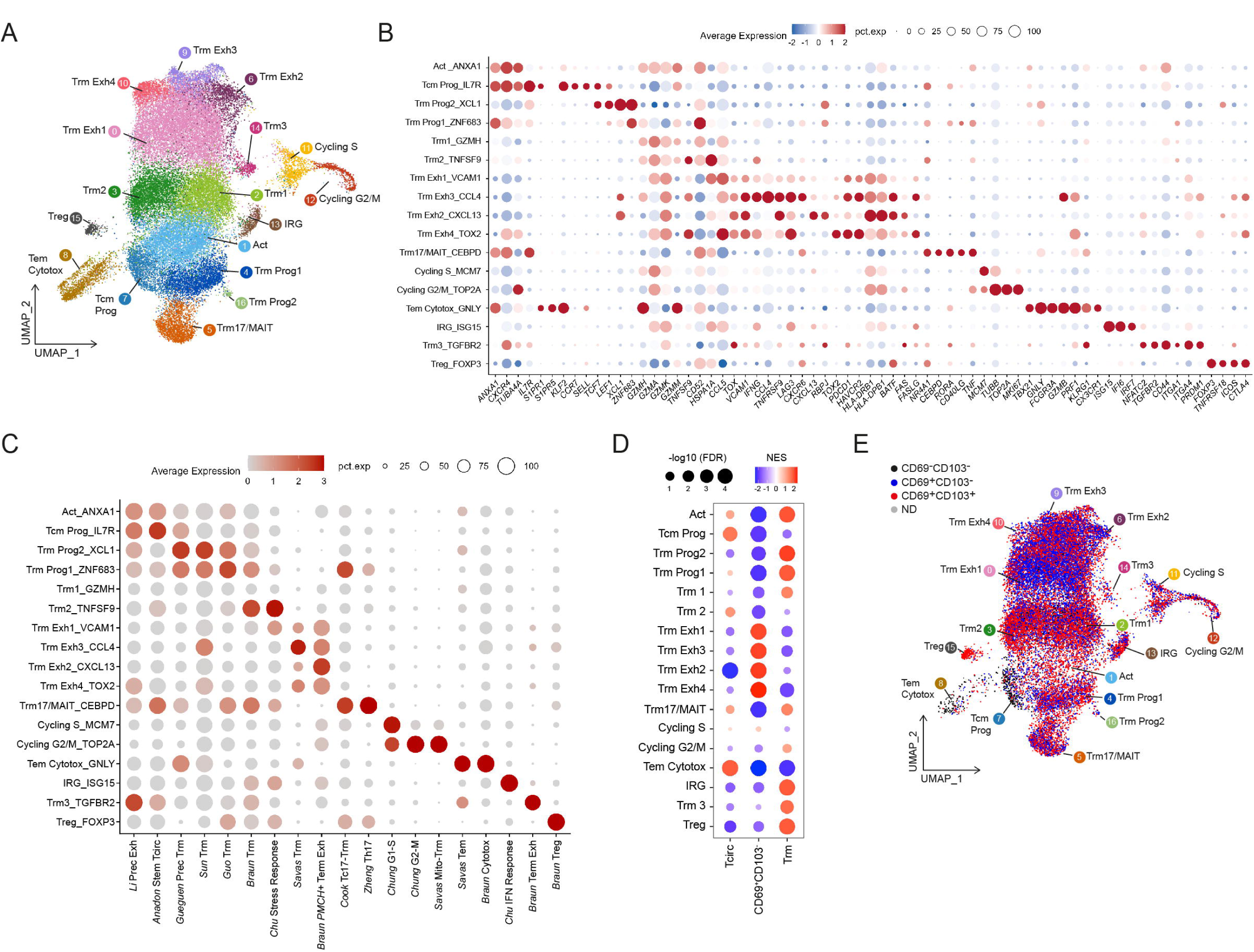
Single-cell transcriptomics reveals high heterogeneity among Trm cells infiltrating ccRCC tumors. Memory CD8^+^ T cells were isolated from ccRCC tumors and analyzed by scRNAseq and CITE-seq **(A)** UMAP visualization of 17 memory CD8^+^ T cell clusters, including a total of 40,721 cells. **(B-C)** Dot plots show key immune-related gene expression **(B)** and expression of curated gene signatures **(C)** across memory CD8^+^ T cell clusters. Dot size indicates the percentage of cells expressing the gene or signature and color intensity indicates average expression. **(D)** Gene Set Enrichment Analysis (GSEA) for scRNAseq clusters gene signatures enriched in Tcirc, CD69^+^CD103^−^ and Trm cells from bulk RNAseq. A signature with positive Normalized Enriched Score (NES) and a -log10-False Discovery Rate (FDR) > 1.3 was considered as positively enriched. A signature with negative NES and a -log10-FDR > 1.3 was considered as negatively enriched. **(E)** UMAP visualization for cell annotation based on TotalSeq CD69 and CD103 protein expression.

We identified two clusters that expressed circulating genes, including *KLF2*, *S1PR1*, and *SELL*. One of these clusters also expressed *CCR7*, progenitor genes such as *TCF7* and *LEF1*, and correlated with the gene signature from stem-like circulating cells, then it was named central memory/progenitor cells (Tcm-Prog). The other circulating cluster expressed genes associated with cytotoxic effector memory T (Tem-Cytotox) cells, such as *FCGR3A*, *GNLY*, *GZMB*, and *PRF1.* This cluster also shared a gene signature described for Tem cells in breast cancer and renal cancer, indicating that it corresponds to cytotoxic Tem cells (Tem Cytotoxic) (**Fig. 4B-C**). All the other clusters exhibited decreased expression of circulating markers (*S1PR1*, *S1PR5*, *KLF2*), indicating tissue retention. Two Trm cell progenitor clusters (Trm Prog1 and Trm Prog2) were characterized by the expression of the transcription factors *ZNF683*, *TCF7* and *LEF1,* and the chemokine *XCL1*. These clusters showed enrichment of the gene signature for Trm progenitors described in non-small cell lung cancer (NSCLC) patients (*Gueguen* Prec Trm) (**Fig. 4B-C**).We identified a cluster that expressed TCR-activation downstream pathways and genes associated with the transition to tissue-resident phenotypes, including *GZMA*, *GZMH* and *GZMK*, indicative of antigen-driven activation (Act). Two clusters, Trm1 and Trm2, also expressed *GZMA*, *GZMH* and *GZMK*, related to cells in transition from progenitors to terminally differentiated or exhausted states^33^. The Trm2 cluster showed high expression of heat-shock protein genes and signatures associated with stress response (*Chu* Stress Response) as well as canonical Trm cells found in ccRCC tumors (*Braun* Trm) (**Fig. 4B-C**). We also found a small cluster of cells expressing the transcription factors *TOX* and *PRDM1*, as well as *TGFBR2* and *ITGA4* and the signatures for precursor exhausted in renal cancer, indicating that these cells are a Trm subset (Trm3) with a precursor exhausted phenotype. Most clusters (Trm Exh1-4) expressed genes associated with T cell exhaustion, like *TOX* and several inhibitory receptors alongside gene signatures for exhausted Trm cells previously described in breast cancer and RCC tumors (**Fig. 4B-C**)^19,35^. Trm Exhausted 1 (Trm Exh1) and Trm Exhausted 2 (Trm Exh2) exhibited lower expression of inhibitory receptors such as *PDCD1* (PD-1), *HAVCR2* (TIM-3), and *LAG-3* compared to Trm Exhausted 3 (Trm Exh3) and Trm Exhausted 4 (Trm Exh4). Trm Exh2 expressed high levels of *CXCL13*, a marker commonly found in tumor-reactive CD8^+^ T cells and associated with favorable responses to ICB^38,39^. Although the Trm Exh3 and Trm Exh4 clusters displayed a terminally differentiated phenotype, they also expressed effector molecules such as *IFNG*, *PRF1* and granzyme B (*GZMB*), indicating their effector potential.

In addition, we identified two clusters of proliferating cells (Cycling S and Cycling G2/M), one cluster enriched in IFN-related genes (IRG), one cluster expressing *RORA*, *CEBPD* and *CD40LG*, all genes associated with IL-17-producing type 3 lymphocytes (Trm17/MAIT), and a cluster of regulatory CD8^+^ T cells (Treg) characterized by the expression of *FOXP3*, *ICOS*, *TNFRSF18* (GITR), and no detectable *CD4* expression (**Fig. 4B-C and Supplementary Fig. 5C**).

We next compared the bulk RNA-seq signatures obtained from the three sorted populations (**Fig. 2**) with the transcriptional profiles of the 17 clusters identified by single-cell RNA-seq. To this end, we selected the top 20 up-regulated genes from each scRNAseq cluster and performed GSEA using bulk RNA-seq samples. As expected, Tcm Prog and Tem Cytotox cluster signatures were enriched in the Tcirc subset, while all exhausted Trm Exh1-4 signatures were enriched in the CD69^+^CD103^−^ Trm-like subset (**Fig. 4D)**. Notably, Trm Prog1-2, Trm17/MAIT, Act, Trm1, Trm3, IRG, and Treg signatures were enriched in the Trm subset (**Fig. 4D)**.

To complement this comparison, we evaluated the protein expression of CD69 and CD103 in one of the patients using antibody-derived tags (ADT) (**Supplementary Fig. 5D**). ADT-based analysis revealed a rather mixed distribution of clusters, with Tcm Prog and Tem Cytotox clusters predominantly enriched in CD69^−^CD103^−^, Trm Prog1 Trm17/MAIT, Act, Trm1, Trm3, IRG, and Treg clusters were enriched in CD69^+^CD103^+^ cells and Trm Exh1-4 clusters enriched within CD69^+^CD103^−^ population (**Fig. 4E**). To further validate these patterns, we grouped cells by CD69 and CD103 expression using ADT, aggregated their transcriptomes, and performed pseudobulk analysis alongside bulk RNA-seq data. PCA demonstrated that pseudobulk samples clustered closely with their corresponding bulk counterparts (**Supplementary Fig. 5E**). Moreover, heatmap visualization of DEGs from the bulk RNA-seq analysis showed highly similar expression profiles whether derived from bulk or pseudobulk data (**Supplementary Fig. 5F**). Together, these results indicate that memory CD8^+^ T cells infiltrating RCC can be classified into three distinct groups: the Tcirc group, comprising Tcm Prog and Tem Cytotox clusters; the CD69^+^CD103^−^ Trm-like group, primarily consisting of exhausted clusters Trm Exh1-4; and the Trm group, which includes rather heterogeneous populations, including Trm Prog1 Trm17/MAIT, Act, Trm1-3, IRG, and Treg clusters.

Our results suggested that memory CD8^+^ T cells infiltrating ccRCC tumors, and particularly Trm cells, represent a more heterogeneous population than previously described. To validate this heterogeneity at the protein level, we analyzed these cells by spectral flow cytometry. We analyzed data from 36,882 CD8^+^ T cells from ccRCC tumor samples from six different patients, performed unsupervised graph-based clustering and identified twelve clusters displayed in a uniform manifold approximation and projection (UMAP) plot (**Fig. 5A**). All the clusters were found in the six ccRCC patients but in different proportions (**Fig. 5B**). We found that most of the CD8^+^ T cells were memory cells, characterized by the expression of CD45RO (**Fig. 5 C-D**), and non-memory cells were mainly effector CD8^+^ T cells (Teff), characterized by the absence of CD45RO and the high expression of cytotoxic markers CX3CR1, granzyme B and granulysin. Memory Tcirc clusters included Tem cells, expressing cytotoxic markers, and Tcm, expressing CCR7 and TCF-1 (**Fig. 5C-D**). Trm cell clusters, expressing both CD69 and CD103, included Trm progenitors (Trm Prog), expressing TCF-1, Trm Exhasuted (Trm Exh) that express PD-1, CD39, intermediated levels of TIM-3 and high levels of Ki67, and Trm expressing only PD-1 (**Fig. 5C-D**). Most of the memory CD8^+^ T cells expressing CD69 but not CD103 (CD69^+^CD103^−^), co-expressed more than one inhibitory receptor PD-1, CD39 or TIM-3, indicating that these cells are terminally exhausted (CD69^+^CD103^−^ Trm Exh1-4). These results closely resemble the heterogeneity found by scRNAseq, validating our findings.

**Figure 5.**
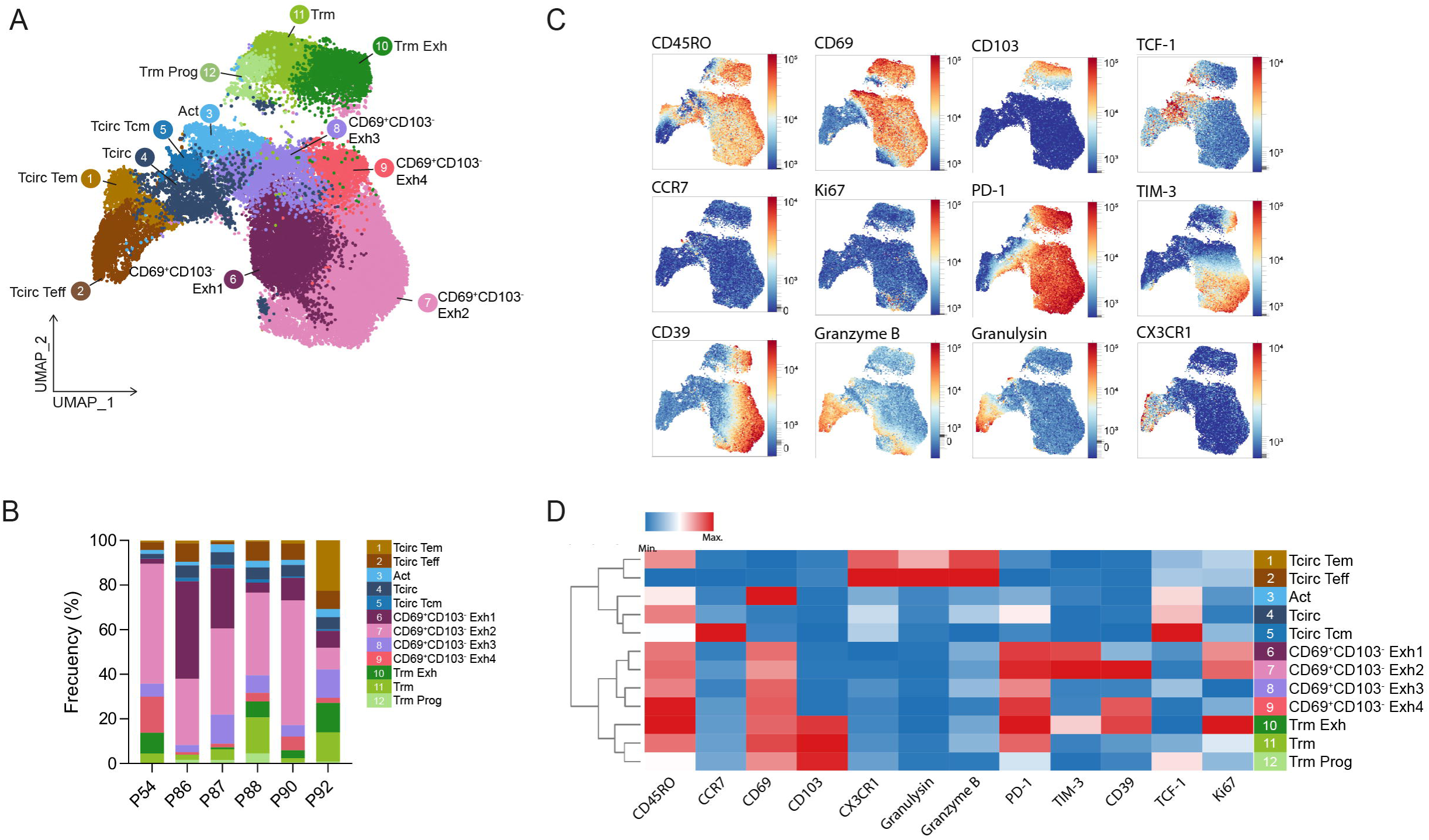
Phenotypic heterogeneity of CD8^+^ T cell infiltrating ccRCC tumors analyzed by spectral flow cytometry. Tumor-infiltrating CD8^+^ T cells from 6 different ccRCC patients were analyzed by spectral flow cytometry. **(A)** UMAP plot showing 12 CD8^+^ T cell clusters identified by FlowSOM (36,882 total cells). **(B)** Frequency quantification of the memory CD8^+^ T cells clusters in each ccRCC patient. **(C)** UMAP plots showing the expression of key markers for cluster annotation. **(D)** Heatmap showing the differential expression of markers used for clustering and annotation.

### TCR and trajectory analyses indicate that central-memory Tcirc cells differentiate into Trm cells that upon clonal expansion progress toward terminal exhaustion

To evaluate the differentiation trajectory of memory CD8^+^ T cells within ccRCC tumors, we performed RNA velocity, PAGA, and pseudotime analyses. RNA velocity revealed a hierarchical flow from Tcm Prog toward Act and Trm-Prog1 states, followed by differentiation into Trm1 and Trm2, with subsequent progression toward exhausted Trm subsets (**Fig. 6A**). Alternative trajectories from Tcm Prog towards Tem Cytotox and Trm17/MAIT suggest additional differentiation branches. Consistently, PAGA identified a strong connectivity backbone linking progenitor-like clusters (Tcm Prog, Trm Prog, Act) to the core Trm network and ultimately to exhausted states, while peripheral populations such as Trm17/MAIT, Treg, and cycling cells showed weaker connections (**Fig. 6B**). For pseudotime estimation, we set the Tcm Prog cluster as the root and ordered cells accordingly (**Supplementary Fig. 6A**). The order of the clusters placed Tcm Prog, Tem Cytotox, Treg, Trm Prog 1-2, and Tc17/MAIT at the beginning; Act and Trm1-2 in the middle; and Trm Exh1-4 and IRG clusters at the end of the pseudotime (**Fig. 6C-D**). Early pseudotime was characterized by high expression of circulating memory and progenitor-associated genes (*KLF2*, *CCR7*, *S1PR1*, *S1PR5*, *TCF7*, *LEF1*), which rapidly declined. Cytotoxic genes showed a temporal shift, with *FCGR3A*, *GNLY*, *GZMH*, *TNF*, and *IL2* expressed early and *GZMB* and *PRF1* increasing at late pseudotime. Tissue-residency programs emerged early, with transient (*ZNF683*, *ITGAE*, *ITGA1*) or sustained (*CXCR6*, *XCL1*, *XCL2*, *CCL4*, *CCL5*, *CCR5*) expression, alongside early Trm17/MAIT-associated genes (*CCL20*, *CCR6*, *CD40LG*, *IL23R*, *RORA*, *CEBPD*). In contrast, exhaustion-associated genes (*HAVCR2*, *PDCD1*, *LAG3*, *TOX*, *TOX2*, *ENTPD1*, *HLA-DRA*, *HLA-DPA1*, *CTLA4*) were progressively upregulated, indicating a continuous trajectory toward exhaustion (Fig. 6E, Supplementary Fig. 6).

**Figure 6.**
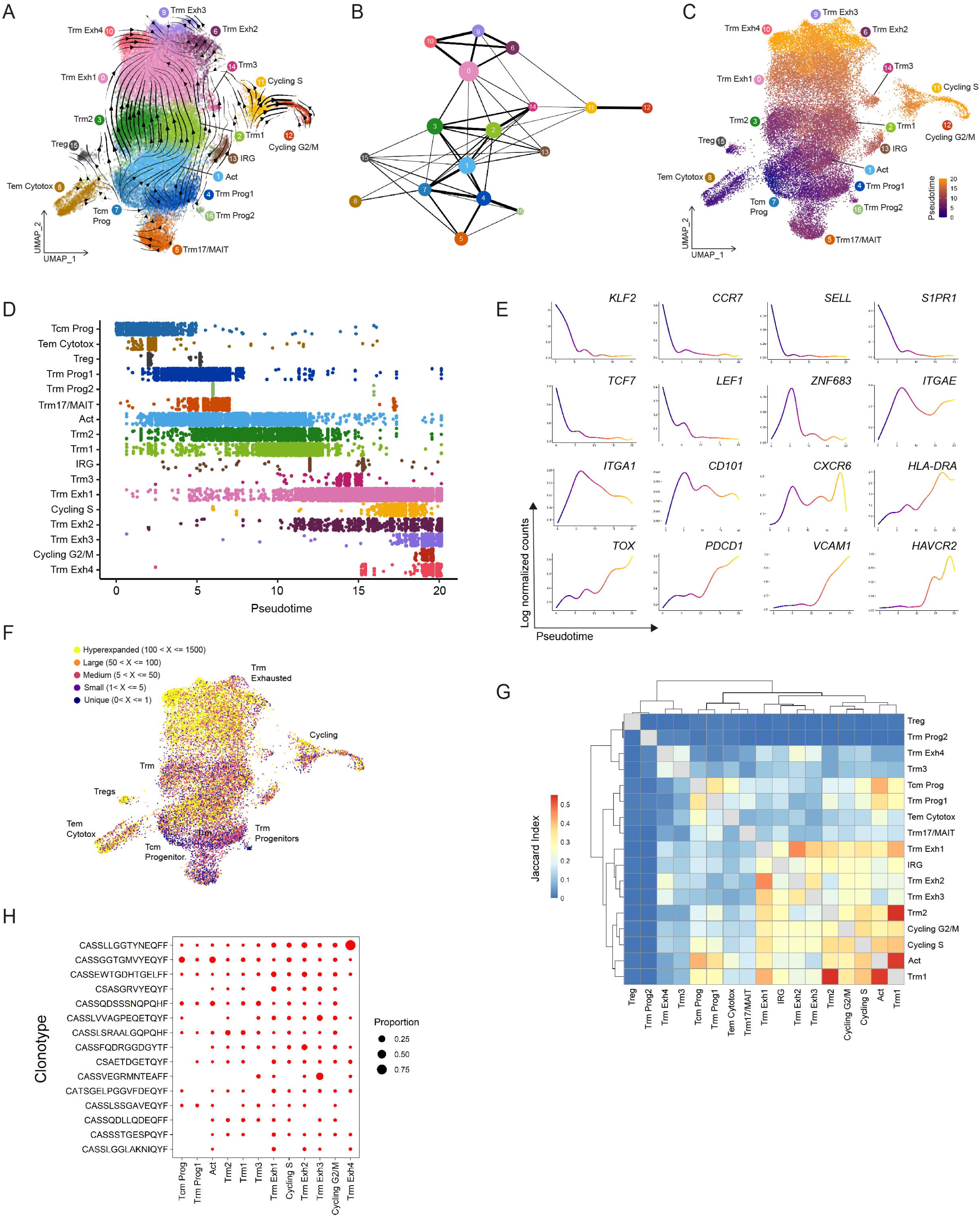
TCR and trajectory analyses indicate that central-memory Tcirc cells differentiate toward Trm cells that progress toward terminal exhaustion. **(A)** UMAP plot displaying RNA velocity vectors of cells from the scRNA-seq dataset. Directional flow of all clusters depicted as streamlines, with line weight increasing with speed. **(B)** Partition-based graph abstraction (PAGA) analysis of memory CD8^+^ T cells clusters from the scRNA-seq dataset. **(C)** UMAP visualization of pseudotime defined by monocle3 trajectory. **(D)** Distribution of memory CD8^+^ TIL clusters across pseudotime. **(E)** Individual curves displaying the gene enrichment of key genes across the pseudotime. **(F)** UMAP visualization of clonotype frequency categorized as hyperexpanded, large, medium, small, and unique. UMAP shows 21,137 cells memory CD8^+^ TILs with TCR data. **(G)** Heatmap showing normalized Jaccard Index of intersected clusters. **(H)** Dot plot representing the distribution of the top 15 expanded clonotypes across main clusters. Dot size indicates the proportion of each clonotype across all clusters. Blank space indicates absence.

To further support the inferred differentiation pathways, we tracked TCR clonotypes across CD8^+^ T-cell clusters using paired RNA-TCR data and TCRβ amino acid sequences. Among 21,137 cells, we identified 3,598 unique clonotypes. Progenitor clusters (Tcm and Trm-Prog1) displayed high clonal diversity and low expansion, yet showed extensive TCR sharing, indicating a common clonal origin among expanded cells (**Fig. 6F, Supplementary Fig. 6C-D**). These progenitor subsets shared even greater clonal overlap with the Act cluster, which exhibited marked clonal expansion, consistent with a TCR-activated state (**Fig. 6G**). In line with the proposed trajectory, Act cells shared the highest fraction of TCRs with Trm1, which in turn showed strong clonal overlap with Trm-Exh1. Stressed-Trm2 cells showed extensive clonotype sharing with Trm1, suggesting a stress-associated state of Trm1 cells. Substantial clonal sharing was also observed among Trm-Exh clusters and between Trm-Exh and Trm1/2, supporting clonal continuity within the tissue-resident compartment (**Fig. 6G**). While Trm1-2 clusters exhibited moderate-to-low expansion, Trm-Exh clusters showed the highest clonal expansion, with hyperexpanded clonotypes (100 < TCRβ ≤ 1,500 cells) predominantly localized to Trm-Exh1–4 and Act-Prog populations. Outside the main differentiation axis, Tem-Cytotox and Treg clusters also contained a substantial fraction of expanded and hyperexpanded clonotypes. Importantly, several expanded clones displayed trajectory-like distributions. For example, the top 15 clonotypes were present at progenitor, cytotoxic, activated, cycling, and exhausted states, albeit at different proportions (**Fig. 6H and Supplementary Fig. 6E**).

In addition, we examined the TCR sharing between ccRCC tumor and NMRT. To do so, first we constructed an integrated NMRT atlas using 5,606 cells from one paired ccRCC-NMRT sample and 14,009 cells from two previously published datasets^19,20^. NMRT data were analyzed using label transfer, UMAP projection, and the integrated ccRCC data as reference. We kept clusters with at least 100 annotated cells. According to label transfer results, the most abundant cluster in NMRT was Tcm Prog, followed by Act Prog, Trm1, Trm2, Trm Prog1, Tem Cytotox, and Trm17/MAIT (**Supplementary Fig. 6F**). Next, we subset the NMRT atlas to keep only our dataset and analyzed TCR data. Hyperexpanded clones were found mostly at Tem Cytotox, Tcm Prog, and Act clusters (**Supplementary Fig. 6G and 6H**). A total of 591 TCRs were shared between RCC and NMRT, representing 6,215 and 1,712 cells, respectively (**Supplementary Fig. 6I**). Next, we filtered-out shared NMRT-RCC TCRs present in less than 10 cells in RCC (kept a total of 107 TCRs). We found that most of these shared TCRs are highly expanded in both tissues (**Supplementary Fig. 6K**). We also found high TCR sharing between NMRT and RCC clusters, particularly in Act, Tcm Prog, Trm Prog1, Trm1, and Trm2 clusters, at both tissues (**Supplementary Fig. 6J**). Taken together, these results indicate that NMRT represents a reservoir of expanded CD8^+^ T cell clones in progenitor memory states.

### Progenitor and activated Trm cell subsets associate with favorable clinical outcomes in ccRCC patients

To evaluate the clinical impact of tumor infiltration of the different memory CD8^+^ T cell subpopulations identified in our study, we estimated their abundance in bulk RNAseq datasets (TCGA-KIRC cohort and the CheckMate 025 trial) through deconvolution using the transcriptional profiles defined by our single-cell analyses and associated with patient survival. In the TCGA-KIRC cohort, we successfully inferred the proportions for nearly all clusters, with the exception of Trm Exh3 and Trm2 (**Supplementary Fig. 7A**). Similarly, in the CheckMate 025 cohort, most clusters were well represented, whereas Cycling G2/M, Trm17/MAIT, Treg, Trm Exh3, Trm Exh1, and Trm1 appeared at lower levels (**Supplementary Fig. 7C**).

Then, survival analyses were performed using Kaplan-Meier curves and log-rank tests analyzing patients divided into two groups according to the optimal cutoff proportion for each subpopulation. Survival analysis revealed opposing clinical outcomes associated with distinct memory CD8^+^ T-cell states. In TCGA-KIRC, patients with high proportions of Trm Prog1, Trm Prog2, Trm3, Tem Cytotox, Trm17/MAIT, Act, and Trm Exh4 clusters exhibited significantly higher overall survival compared (**Fig. 7A**). In contrast, enrichment of Trm1, Treg, IRG, Cycling, and Trm Exh2 signatures was associated with worse survival (**Fig. 7B**). Other clusters showed no significant association with patient outcomes (**Supplementary Fig. 7B**). A similar pattern was observed in the CheckMate 025 cohort. Patients with high proportions of Trm Prog1 and Act clusters survived longer (**Fig. 7C**), whereas those with elevated Cycling G2/M, IRG, and Trm Exh2 signatures had shorter survival (**Fig. 7D**). No significant associations were detected for the other subsets in this cohort (**Supplementary Fig. 7D**). Together, these results suggest that distinct CD8^+^ T cell states have divergent clinical associations in ccRCC, with progenitor-like and activated Trm subsets conferring favorable outcomes, whereas exhausted, regulatory, and cycling states mark more aggressive disease.

**Figure 7.**
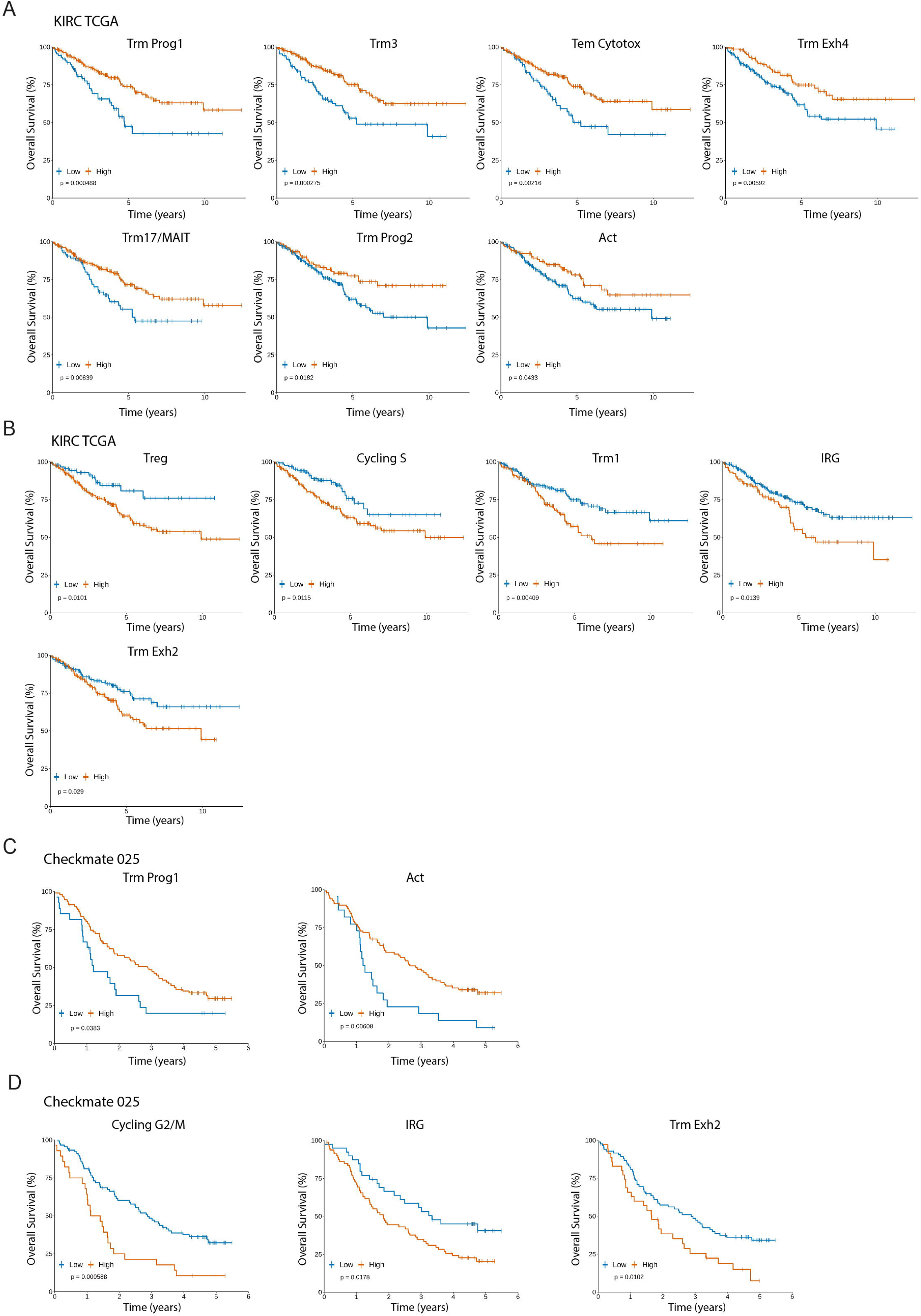
Distinct memory CD8^+^ T cell subsets associate with opposing clinical outcomes in ccRCC patients. Single-cell expression matrix was used to deconvolute The Cancer Genome Atlas- Kidney Renal Cell Carcinoma (TCGA-KIRC) and Checkmate-025 datasets and survival analysis was performed for each cluster. Kaplan-Meier plots of the clusters associated with **(A)** longer survival and **(B)** poor survival at TCGA-KIRC dataset. Kaplan-Meier plots of the clusters associated with **(C)** longer survival and **(D)** poor survival at Checkmate-025 dataset. Patients were stratified into ‘High’ (red) and ‘Low’ (blue) abundance groups for each immune variable using an optimal cut-point, determined by the maximally selected rank statistic. Log-rank test was used to estimate p values.

## Discussion

In contrast to most cancers, a higher density of tumor-infiltrating CD8^+^ T cells in ccRCC associates with poorer clinical outcomes^40^. Our findings provide a mechanistic explanation for this paradox by showing that tumor-infiltrating CD8^+^ T cells acquire functionally heterogeneous differentiation states with opposing clinical associations in ccRCC. We show that Trm cells exhibiting high tumor reactivity against autologous ccRCC cells, adopt more functional states, including progenitor-like Trm cells, which correlate with favorable survival and improved responses to immunotherapy, which is consistent with observations in other malignancies such as ovarian cancer, melanoma and head-and-neck carcinoma^35,41^. Moreover, the majority of tumor-infiltrating CD8^+^ T cells display an exhausted or terminally exhausted phenotype, consistent with previous reports showing poor clinical outcomes of overall CD8^+^ T cell infiltration^19,20^. Hence, the differentiation and functional landscape of tumor-resident CD8^+^ T cells rather than total amounts, determines the clinical outcomes infiltration in ccRCC. Multidimensional analyses centered on CD8^+^ T cell populations revealed a substantial heterogeneity in the tissue resident compartment, including progenitor, activated, transition, and exhausted Trm subsets. The differentiation pathways among memory and exhausted CD8^+^ T cells within human tumors remain debated^42^. Our data favor a linear differentiation continuum in which T cells switch from a circulating program with higher stemness and cytotoxic potential towards tissue-resident profiles that transition into exhausted states. Trajectory and TCR-sharing analyses suggest that circulating progenitors (Tcm Prog) give rise to tissue-resident progenitors (Trm Prog1-2) and both subsets converge into a TCR-activated state (Act) characterized by higher clonal expansion and upregulation of effector molecules *GZMH*, *GZMA* and *GZMK*, that progressively transition towards exhausted Trm subsets. This supports a progressive model of Trm differentiation, bridging circulating precursors to terminally differentiated tumor-resident cells. Similar trajectories have been observed where circulating precursors transition through intermediate Trm cell states before acquiring full exhaustion. As an example, a study in ovarian cancer showed that recirculating progenitor CD8^+^ T cells expressing TCF7, differentiate into progenitor Trm cells that transition into effector and exhausted states^35^. In lung cancer, circulating (KLF2^+^) or resident (XCL1^+^) progenitors differentiate first into a transitional state characterized by the expression of GZMH and terminally differentiate into exhausted cells with resident phenotype^33^. In our data, the effector molecule GZMH is expressed by Trm-Prog1 and transitional Act and Trm clusters and then downregulated in Trm-Exh clusters, whereas GZMK marks transitional Act and Trm clusters and also Trm-Exh clusters. Together, these findings reveal that acquisition of a tissue-resident program precedes the onset of exhaustion, positioning Trm differentiation as an preceding step bridging progenitor to terminally exhausted CD8^+^ T cell states within tumors.

In contrast to circulating subsets, particularly Tem cells, that retain robust expression of cytotoxic mediators in the tumors, the tissue-resident compartment showed reduced expression of cytotoxic molecules such as perforin and granzyme B, which largely mediate elimination of tumor targets^43^. This observation is consistent with recent evidence showing that the cytotoxic program in human memory CD8^+^ T cells is suppressed by the acquisition of a tissue residency program driven by environmental cues, such as IL-15 and TGF-β^44^. Similarly, circulating NK cells entering solid tumors are reprogrammed within a day into CD49a^+^ tissue-resident cells that lose killing ability and stop recruiting other immune cells^45^. Nevertheless, we observed that after *in vitro* expansion, Trm cells can regain antigen-specific production of granzyme B and perforin, indicating that this repression may be reversible. Furthermore, both Trm and CD69^+^CD103^−^ Trm-like subsets expressed FASLG, IFNγ, TNFα, *GZMH*, *GZMA* and *GZMK* suggesting that these populations may mediate antitumor activity through noncanonical cytotoxic pathways. In particular, it has been recently reported that CD8^+^ T cells can produce granzyme K and mediate the activation of the complement cascade^46,47^. In addition, CD8^+^CD40L^+^ T cells in ccRCC induce tumor cell apoptosis through CD40-dependent caspase-8 activation, independently of perforin and granzyme-mediated cytotoxicity, and a CD40L resistance signature associates with inferior survival in TCGA-KIRC^48^. Even if Tem cells have low TCR sharing with clones following the main trajectory pathway, it has been shown that the abundance of peripheral T cells exhibiting a cytotoxic profile and sharing TCR clonotypes with tumor-infiltrating T cells correlates with better responses to anti-PD-L1 therapy^49^, supporting the relevance of these cytotoxic populations.

In addition to the Trm subsets involved in the main differentiation trajectory, other subsets also emerged in our analysis, reflecting functional diversity of memory CD8^+^ T cells. Trm cells with a Th17/Tc17 signature have been shown to contribute to antitumor immunity by maintaining polyfunctionality and tissue persistence^50^. High levels of IFNγ^−^ Tc17 CD8^+^ T cells have been associated with poor survival in hepatocellular carcinoma^51^. In ccRCC, however, a Th17-derived signature correlates with improved outcomes in TCGA-KIRC and is the only immune-related signature linked to better survival^40^. ccRCC tumors produce high levels of IL-6, a cytokine that promotes Th17/Tc17 differentiation, which may support the emergence of Trm17/MAIT cells^52^. MAIT cells in healthy kidney tissue also show a resident phenotype and produce IL-2, GM-CSF, and IL-17A, indicating functional relevance^53^. Consistently, we found that a high proportion of the Trm17/MAIT cluster is associated with longer survival. Th17/Tc17 cells can exert antitumor activity through their plasticity toward Th1/Tc1-like cytotoxic states and by promoting dendritic cell recruitment and cross-priming^54,55^. Conversely, we also identified regulatory CD8^+^ T (Treg) cells in ccRCC tumors, a population with limited prior characterization. In colorectal cancer, CD8^+^ Treg cells inhibit proliferation and cytokine production by Th1 cells^56^. In prostate cancer, CD8^+^ Treg cells have been shown to inhibit proliferation of naïve T cells, and their suppressive function could be reversed by TLR signaling^57^. The expression of CD103 in Trm cells is induced by TGF-β, allowing Trm to permanently reside in the tissue^58^. TGF-β is also important during generation of CD8^+^FOXP3^+^ Treg and Tc17 cells, and CD103 is expressed in both cell types in a context-specific manner^59^. Thus, taking into consideration that TGF-β is a key driver of tumorigenesis and metastasis, it is not surprising to find CD8^+^ Treg and Tc17 subpopulations alongside Trm in ccRCC tumors^60^.

In summary, our integrative single-cell and functional analyses reveal that ccRCC harbors a continuous spectrum of tumor-specific Trm cells, encompassing progenitor, transition, and terminally exhausted subsets correlating with opposing clinical outcomes. These findings reconcile the long-standing paradox of immune infiltration in ccRCC, by showing that it is not the abundance of CD8^+^ T cells per se that determines clinical outcome, but rather their functional states, which ultimately define the efficacy of the antitumor immune response.

## Methods

### Human samples

Freshly surgically resected clear cell renal cancer (ccRCC) tumor tissue, non-malignant renal tissue, and whole blood were collected at Hospital Barros Luco (Santiago, Chile) under a dedicated protocol approved by the Scientific Ethical Committee of the Servicio de Salud Metropolitano Sur (memorandum N°318/2017) following written informed consent. This study considered patients who underwent partial or radical nephrectomy for tumor removal (stage I to III) without prior systemic treatment. Tumor and non-malignant renal tissue samples from RCC patients treated at the Foch Hospital (Suresnes, France) were analyzed using single-cell sequencing (scRNAseq) at the Institut Curie (Paris, France). Both biological material and patients’ personal data were safeguarded and regulated under the Material Transfer Agreement (MTA) n°20190052 and the regulation (EU) 2016/679 of the European Parliament.

### Tissue processing

All samples were processed within 4 hours after surgery. Fresh ccRCC tumors and non-malignant renal tissue samples were cut into small pieces in enzymatic digestion media (RPMI-1640 (HyClone), 0.5% Collagenase I-II-IV (Gibco), 0.025% DNase (Roche), 0.1% Dispase (Gibco), 0.25% Hyaluronidase (Sigma-Aldrich), and 1% penicillin-streptomycin (Corning)). Samples were incubated for 15-20 minutes at 37°C for digestion, and then the enzymatic reaction was stopped with supplemented RPMI medium (RPMI-1640 (HyClone), 10% fetal bovine serum (SBF, Gibco), 1% penicillin-streptomycin (Corning), 1% non-essential amino acids (NEM, Gibco), 1% Glutamax (Gibco), 1% Pyruvate (Gibco). Dissociated cells were filtered using a 70 µm cell strainer. Cells were resuspended in RBC (BioLegend) to lyse red blood cells and resuspended in supplemented RPMI (for flow cytometry and functional assays) or CO_2_-independent medium (Gibco) with 0.4% human albumin (Vialebex) (for transcriptomics analysis). PBMC from fresh whole peripheral blood were isolated by density gradient centrifugation using Lymphoprep (Stemcell Technologies) according to the manufacturer’s instructions and resuspended in supplemented RPMI.

### Flow cytometry

From 0.5 to 1 million cells were aliquoted into a 96-well U-bottom plate. Cells were incubated with a mixture of Human TruStain FcX (BioLegend) and the viability dye Zombie Aqua or Zombie Near Infrared (BioLegend) for 20 minutes at 4°C in the dark. Chemokine receptor antibodies were labeled in a staining buffer (50% FACS Buffer (PBS with 2% FBS) and 50% Brilliant Stain Buffer (BD Bioscience)) for 20 minutes at 37°C protected from light. Cell surface labeling was performed in staining buffer for 30 min at 4°C in the dark. Cells were fixed and permeabilized for 40 minutes at room temperature, protected from light, and stained for intracellular or intranuclear targets using eBioscience FOXP3 Transcription Factor Staining Buffer Set (Invitrogen) for 40 minutes at room temperature, also protected from light. Once marked, cells were resuspended in FACS buffer and acquired using a FACSymphony A1 (BD Bioscience) and Aurora (Cytek) cytometers. Data were analyzed using FlowJo^TM^ v10.10 software (BD Bioscience) and OMIQ (Dotmatics).

### Cell sorting

Cells from ccRCC tumor were incubated with a mixture of Human TruStain FcX (BioLegend) and the viability dye Zombie Aqua (Biolegend) for 20 minutes at 4°C in the dark followed by cell surface labeling using anti-CD4, anti-CD8, anti-CD45RO, anti-CD69 and anti-CD103 antibodies to identify and isolate single-cell live memory CD8^+^ T cells subpopulations sorted as: Tcirc cells (Zombie Aqua^−^CD4^−^CD8^+^CD45RO^+^CD69^−^CD103^−^); CD69^+^CD103^−^ cells Zombie Aqua^−^CD4^−^CD8^+^CD45RO^+^CD69^+^CD103^−^); Trm cells (Zombie Aqua^−^CD4^−^CD8^+^CD45RO^+^CD69^+^CD103^+^). Αnti-CD3 antibody was not used to avoid cell activation. Sorting was done using a BD FACSAria II cell sorter (BD Biosciences). For functional assays, sorted cells were in vitro expanded using a human CD8^+^ T cell expansion protocol. For bulk RNAseq, cells were resuspended in Buffer RLT lysis buffer (Qiagen) and frozen at −80°C until use. For scRNA-seq and TCR-seq, sorted cells were loaded into a 10X Chromium instrument (10X Genomics) within 6 hours. For patient RCC03, cells were first marked with TotalSeq antibodies for CD69 and CD103 (BioLegend) before being loaded into the 10X instrument.

### In vitro expansion of human CD8^+^ TILs

Sorted memory CD8^+^ T cells subsets were resuspended in T cell expansion medium (TexMACS medium (Miltenyi Biotec), 1% penicillin-streptomycin (Corning), 1% non-essential amino acids (NEM, Gibco), 1% Glutamax (Gibco), 1% Pyruvate (Gibco), 10mM HEPES (Gibco), 50 µM 2-mercaptoethanol (Gibco)), supplemented with 30 ng/mL recombinant human anti-CD3 (OKT3 clone, Biolegend) and 6000 IU/mL recombinant human IL-2 (Proleukin, Novartis) at a concentration of 0.5-1×10^6^ cells/mL and were incubated at 37°C and 5% CO_2_. After 48h, cells were washed, resuspended in T cell expansion medium with 6000 IU/mL IL-2, which was replaced every 3-5 days until they were used for the tumor cell recognition assay.

### Generation of primary cultures of ccRCC cells and non-malignant renal tissue

1-2×10^6^ cells obtained from ccRCC tumors and nonmalignant renal tissue single-cell suspensions were seeded on T75 cell culture flasks coated with collagen I (Thermo Scientific) in Renal TumorMACS medium (Miltenyi Biotec) for ccRCC cells and supplemented RMPI for NMRT and incubated at 37°C and 5% CO_2_ overnight. The next day, the T75 flasks were washed twice with PBS to remove non-adherent cells. Cells were maintained in culture for two weeks and kept at 70% confluency. To verify the purity of the primary cultures, EpCAM expression was measured by flow cytometry, and those cultures with more than 95% EpCAM^+^ cells were considered pure epithelial cultures.

### Autologous tumor cell recognition assay

Recognition of autologous ccRCC tumor cells by expanded memory CD8^+^ T cells was assessed by measuring increased 4-1BB expression as well as IFN-γ and TNF-α release in coculture assays^41^. The co-culture experiment was performed after 14 to 21 days of CD8^+^ T cell expansion. 1×10^5^ expanded CD8^+^ T cells were cultured with 1×10^5^ autologous tumor cells in supplemented RPMI medium without IL-2. As a control for the specificity of antigen recognition presented by the tumor cells, tumor cells were preincubated with 50 µg/mL anti-HLA class I blocking antibody (clone W6/32, BioLegend) for 1 h before addition of the T cells. To confirm specific reactivity against tumor antigens and not self-antigens, T cells were cultured with non-malignant renal tissue cells. As a control for basal expression of 4-1BB, 1×10^5^ expanded T cells were cultured in medium supplemented without IL-2. After 16 h, supernatants were collected and IFN-γ and TNF-α levels were measured by CBA (Th1/Th1/Th17 Human CBA kit, BD Bioscience) and granzyme B and perforin levels were measured by LegendPlex (Biolegend). Cells were harvested and stained for 4-1BB overexpression by flow cytometry.

### Bulk RNA-seq library preparation

Sorted memory CD8^+^ populations were collected and lysed in RLT lysis buffer (Qiagen). RNA was isolated with the Single Cell RNA Purification Kit (Norgen). RNA integrity was assessed with an RNA 6000 pico kit (Agilent). Libraries were generated with the Nextera XT Preparation kit and sequenced on Illumina NovaSeq X using 100 bp paired-end mode and 25 million reads per sample.

### Bulk RNA-seq data analysis

FASTQ files were trimmed with TrimGalore! and aligned against GRCh38A/hg38 with HISAT2. Reads per gene were counted with featureCounts. Raw count matrix was imported into R (v4.2.2) and analyzed with DESeq2 (v1.38.3). Genes with less than 50 counts across all samples were removed. Data was normalized using the variance stabilizing transformation (vst) method. Patient batch effect was removed from the analysis by adding to DESeq2 formula and from the vst count matrix with limma (v3.54.2). Differential expression analysis was conducted with negative binomial regression and Wald’s test of significance. A gene was considered as a differentially expressed gene (DEG) when the adjusted p-value < 0.05.

### Gene Set Enrichment Analysis

Vst-transformed and -batch corrected count matrix was imported into Gene Set Enrichment Analysis (GSEA) tool (v4.3.2). CD8 gene signatures from public data were manually curated by retaining only the top 20 upregulated genes as determined by fold change. A signature was considered as enriched when the adjusted p-value < 0.05.

### Single-cell RNA and TCR-seq library preparation

Memory CD8^+^ T cells were sorted from ccRCC and non-malignant renal tissue samples and loaded into 10X Chromium (10X Genomics). Libraries were prepared using Single-cell 5’ V(D)J kit (Immunoprofiling kit v.1.1, 10X Genomics) according to the manufacturer’s protocol. 5’GEX libraries were assessed for quality and sequenced using the Illumina NovaSeq 6000 with paired-end mode 26 x 98 bp and targeting at least 50,000 reads per cell. After the generation of 5’GEX libraries, V(D)J segments were enriched to construct TCR libraries. Quality of V(D)J libraries was sequenced with Illumina NovaSeq X using paired-end 150 bp.

### Single-cell RNA-seq data analysis

Base call files (BCL) were demultiplexed and converted into FASTQ files with Bcl2fastq2 (v2.20). Cellranger count (v3.1.0) was run on each FASTQ file and GRCh38/hg38 was used as a reference genome. Files produced by cellranger were loaded into R and processed and analyzed with Seurat (v4.3.0.1)^61^. Cells with < 200 and > 2,500 expressed genes, and > 5 % mitochondrial genes, were removed to filter debris and dead cells. Doublets were estimated and filtered with DoubletFinder (v2.0.3). Genes expressed in less than 3 cells, TCR genes, and *MALAT1* were also filtered. Clusters with less than 25 % of cells with paired RNA-TCR data were removed. Data was normalized using the NormalizeData function and LogNormalize method. 2,000 most variable genes were estimated using FindVariableFeatures function and vst method. Low and high cutoffs for feature dispersion and mean were fixed at 0.5 to Inf and - Inf to Inf, respectively. To integrate three ccRCC samples, anchor finding and canonical correlation analysis with the top 30 principal components (PCs) were applied. After integration, the data was scaled with the ScaleData function and linear transformation. Principal component analysis was applied to integrated data and PCs were visualized in an Elbow plot. The top 30 PCs were used to construct a shared nearest-neighbor graph, and clusterization was determined with the Louvain algorithm and visualized using Clustree (v0.5). To visualize data, uniform manifold approximation and projection (UMAP) dimension reduction was applied to the top 30 PCs. After the first round of integration, two clusters with low TCR data paired to RNA were eliminated, and a second round of integration was then performed as indicated above. A total of 40,742 cells were kept after quality control and integration steps. To determine DEGs between clusters, the FindAllMarkers function and the Wilcoxon rank sum test were used with a minimum log_2_ fold-change (log_2_FC) of 0.25, a minimum fraction of 0.25 of cells expressing the gene, and an adjusted p-value < 0.05. For signature analysis, manually curated CD8 signatures were analyzed with the AddModuleScore function. For cluster annotation, two steps were taken: (1) Manual inspection of the up- and down-regulated DEGs and (2) CD8 signature enrichment analysis per cluster.

### Label transfer and projection onto integrated reference

An annotated dataset of three integrated RCC samples was used as a reference for cell type classification of non-malignant renal tissue (NMRT) from one patient of our ccRCC dataset and from two previously published datasets^19,20^. Quality control of the NMRT dataset was assessed as mentioned above. Anchors between reference and NMRT datasets were found and transferred using FindTransferAnchors and TransferData with the top 30 PCs, respectively. An annotated NMRT dataset was projected onto the integrated ccRCC reference UMAP using the MapQuery function, “pca” as reduction reference, and “umap” as reduction model.

### CITE-seq data analysis

Antibody-derived tags (ADTs) against CD69 and CD103 were analyzed with Seurat. Protein expressions were normalized using the CLR method and cells were annotated as double positive (CD69^+^CD103^+^), single positive (CD69^+^CD103^−^), or double negative (CD69^−^CD103^−^) depending on the expression of CD69 and CD103.

### Velocity and trajectory analysis

Seurat object containing annotated clusters was converted into anndata format and imported into Python (v3.9.13). Cellranger output was also imported into Pyhton and spliced and unspliced count matrix were constructed using velocyto (v0.17). The resulting loom files were merged with an anndata file containing cell embeddings and cluster annotations. RNA velocities were estimated with scVelo (v0.2.5) using the stochastic model of transcriptional dynamics. To explore data topology and cluster connectivity, a graph-like map was generated with partition-based graph abstraction (PAGA) using Scanpy (v1.9.1). For trajectory analysis, Seurat object containing cluster annotations and cell embeddings was converted into cell dataset format and analyzed with Monocle3 (v1.3.1). A single principal graph was learned with learn_graph function and a minimal_branch_len of 20. To order cells and estimate pseudotime, cluster “Tcm Prog” was set as root. DEGs across pseudotime were calculated with graph_test function. A gene was considered as DEG when Moran’s I > 0.1 and adjusted p value < 0.05.

### Single-cell TCR-seq data analysis

FASTQ files were processed using cellranger vdj function and GRCh38/hg38 as reference genome. The generated filtered counting files were imported into R and analyzed using scRepertoire (v.1.11)^62^. CombineTCR function was used to filter barcodes with more than two chains and with no value for at least one chain. Cells with paired TRA and TRB chains were considered as clones, and only TRB amino acid information was used for downstream analysis. The level of clonotype expansion was defined as follows: 2 – 5 = “small”, 6 – 50 = “medium”, 51 – 100 = “Large”, 100 < TCRβ ≤ 1,500 = “hyperexpanded”. combineExpression function was used to merge TCR and RNA data. A total of 21,137 cells with paired RNA-TCR data were recovered. For diversity analysis Inverse Simpson Index was used and estimation of shared clonotypes between clusters was calculated using Jaccard index. To visualize TCR sharing between clusters as a chord diagram, first we use getCirclize function to estimate the clone sharing across clusters and circlize (v.0.4.15) for plotting.

### Deconvolution of TCGA-KIRC RNA-seq data

Clinical, gene expression (TPM), and survival data from the TCGA Kidney Renal Clear Cell Carcinoma (TCGA-KIRC) cohort were retrieved using TCGAbiolinks. Overall survival (OS) was calculated from initial diagnosis to death (any cause), censoring living patients at the last follow-up. Cell-type fractions were estimated using CIBERSORTx with a custom signature matrix derived from our scRNA-seq dataset, ensuring a direct linkage to the 17 memory CD8^+^ T cell clusters identified in our study^63^. The signature matrix was generated with the following parameters: 300–500 genes per cell type, 50% random cell sampling with 100 replicates, and gene inclusion requiring expression in ≥80% of cells per phenotype. S-mode batch correction was applied to reduce technical biases between single-cell and bulk RNA-seq datasets. The resulting batch-corrected fractions were used for downstream survival analyses. Deconvolution results were merged with clinical annotations, and overall survival (OS) was used as the primary endpoint.

### Checkmate-025 bulk deconvolution and survival analysis

Bulk expression profiles from the CheckMate 025 trial were deconvolved using the same scRNA-seq–derived reference, employing the full set of 17 memory CD8^+^ T cell cluster signatures and the same CIBERSORTx Fractions workflow with S-mode batch correction enabled. Deconvolution results were merged with clinical metadata by patient identifier. Overall survival time was obtained from the study-provided OS variable (months) and converted to days; the corresponding censoring indicator was used to define the event variable according to the dataset coding. Associations between inferred cell-state fractions and overall survival were evaluated using the same univariate survival framework applied to the TCGA-KIRC cohort.

### Univariate Survival Analysis and Optimal Cut-point Determination

For each inferred memory CD8^+^ T cell cluster fraction, univariate survival analyses were performed using Kaplan–Meier estimation with log-rank testing and Cox proportional hazards regression. For visualization and stratified comparisons, continuous fractions were dichotomized using maximally selected rank statistics (maxstat), selecting the cut-point that maximizes the log-rank statistic across candidate thresholds. To minimize instability and prevent extreme stratification, candidate cut-points were constrained to exclude thresholds assigning fewer than 20% or more than 80% of patients to either group. P-values derived from maximally selected statistics were computed using the standard correction for cut-point selection.

## Acknowledgments

We thank Maria José Fuenzalida, Director of the Cytometer Facilities at Fundación Ciencia & Vida for her technical assistance in cell sorting. We would like to acknowledge the Institut Curie’s CurieCoreTech platforms for their technical and scientific expertises, and Coralie Guérin, Léa Guyonnet, Anne-Gaëlle Lafont, Anna Chipont, Annick Viguier, at Curie’s Cytometry platform (CYTPIC). High-throughput sequencing was performed by the ICGex NGS platform of the Institut Curie supported by the grants ANR-10-EQPX-03 (Equipex) and ANR-10-INBS-09-08 (France Génomique Consortium) from the Agence Nationale de la Recherche (“Investissements d’Avenir” program), by the ITMO-Cancer Aviesan (Plan Cancer III) and by the SiRIC-Curie program (SiRIC Grant INCa-DGOS-465 and INCa-DGOSInser12554). Data management, and quality control were performed by the Bioinformatics platform of the Institut Curie. We also thank Sonia Lameiras, Virginie Raynal, Benoit Albaud, and Patricia Legoix from NGS Platform.

## Funding

This work was funded by Centro Basal Ciencia & Vida, FB210008, FONDECYT grants 1212070 and 1251312 (A.L), 11190897 (V.B), 11261520 (A.H.O), 3220788 (S.H), 3220741 (A.H.O), 3230650 (F.S) 3240458 (D.F) and 3260791 (S.H.G), FONDEQUIP EQM220027 from Agencia Nacional de Investigación y Desarrollo (ANID); Fondation pour la Recherche Médicale (FRM, grant number EQU202203014661), the “Agence Nationale de la recherche” (ANR-15-C E13-009), the LabEx DCBIOL (ANR-10-IDEX-0001-02 PSL; ANR-11-LABX-0043); and Center of Clinical Investigation (CIC IGR-Curie 1428). PSL Global Seed Fund – Ref. PSL: 2023-168, INCa-DGOS-Inserm 12554.

## Author contributions

Conceptualization: S.H, A.H.O, V.B.C, E.P, A.L

Methodology: S.H, A.H.O, J.T.B, E.L, Y.M.K, W.R, C.B.

Investigation: S.H, A.H.O, J.T.B, J.R.A, E.L, C.B, F.S, M.F, E.R, D.F, T.L, C.R, Y.A.

Formal Analysis: S.H, A.H.O, J.T.B, J.R.A, C.B, S.H.G, V.B.

Project Supervision: S.H, A.H.O, V.B.C, E.P, A.L

Funding acquisition: S.H, A.H.O, F.S, D.F, S.H.G, D.S, F.G.C, V.B.C, E.P, A.L

Writing – original draft: S.H, A.H.O, E.P, A.L.

## Competing interests

E.P. is co-founder for Egle Therapeutics. The remaining authors declare no competing interests.

## Data availability

Raw and processed bulk RNA-seq was deposited at Sequence Read Archive and Gene Expression Omnibus under accession number GSE318831. Raw and processed single-cell RNA-seq, single-cell TCR-seq, and single-cell CITE-seq data are available under accession number GSE318834.

## Supplementary Figures legends

**Supplementary Figure 1.**
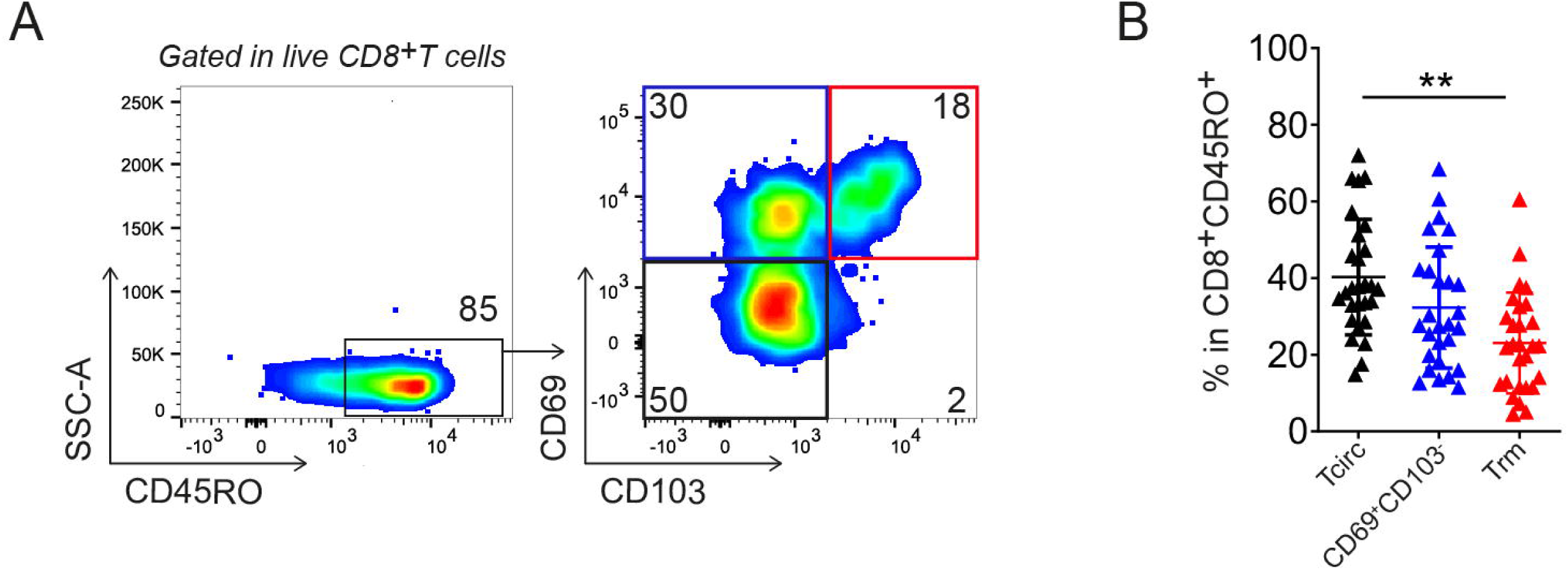
Frequencies of memory CD8^+^ T cell subpopulations in non-malignant renal tissue. **(A)** FACS plot showing the expression of CD69 and CD103 in memory CD8^+^ T cells from non-malignant renal tissue (NMRT). **(B)** Frequency quantification of memory CD8^+^ T cells subpopulations presents in NMRT samples from RCC patients. Each point represents a patient. Statistical analysis was performed using Friedman paired one-way ANOVA test. *p < 0.05; **p < 0.01; ***p < 0.001; **** p < 0.0001.

**Supplementary Figure 2.**
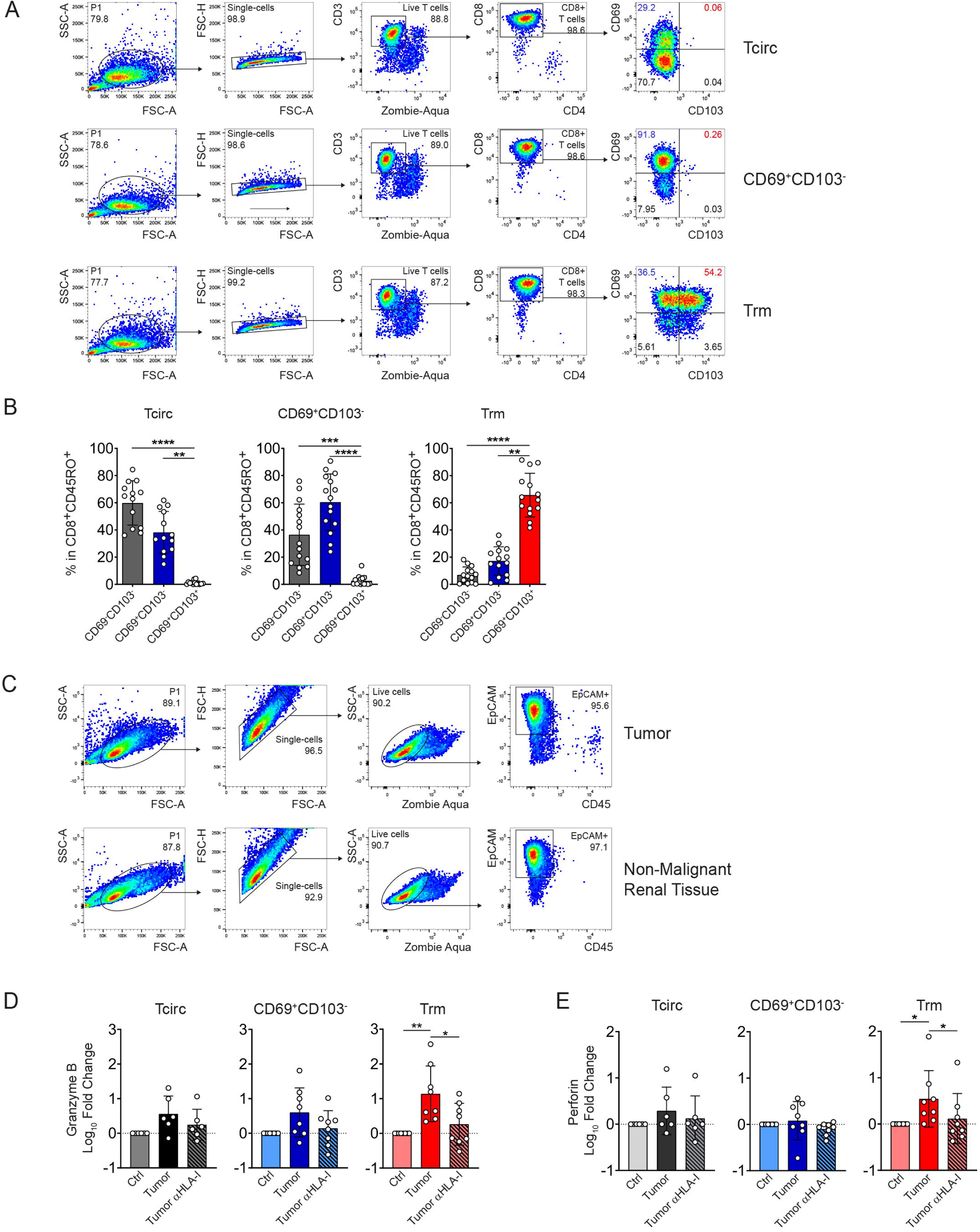
Phenotype of *ex vivo* expanded memory CD8+ T cell subsets, cRCC and NMRT cells and production of cytotoxic molecules after coculture assay. **(A)** FACS plots showing the gating strategy for the analysis of CD69 and CD103 expression in *in-vitro* expanded memory CD8^+^ T cell subpopulation. **(B)** Quantification of change in Tcirc, CD69^+^CD103^−^ and Trm frequency population in *in-vitro* expanded memory CD8^+^ TILs subpopulation cells. **(C)** Representative FACS plots showing the gating strategy for the analysis of EpCAM expression in primary cultures for ccRCC cells and non-malignant renal tissue. **(D-E)** Granzyme B **(D)** and Perforin **(E)** concentrations were determined for each co-culture condition in Tcirc, CD69^+^CD103^−^ and Trm cells. Log_10_ Fold change values were calculated using Granzyme B and perforin concentrations in only T cell condition (Ctrl). Each point represents a patient. Statistical analysis was performed using Friedman paired one-way ANOVA test. *p < 0.05; **p < 0.01.

**Supplementary Fig. 3.**
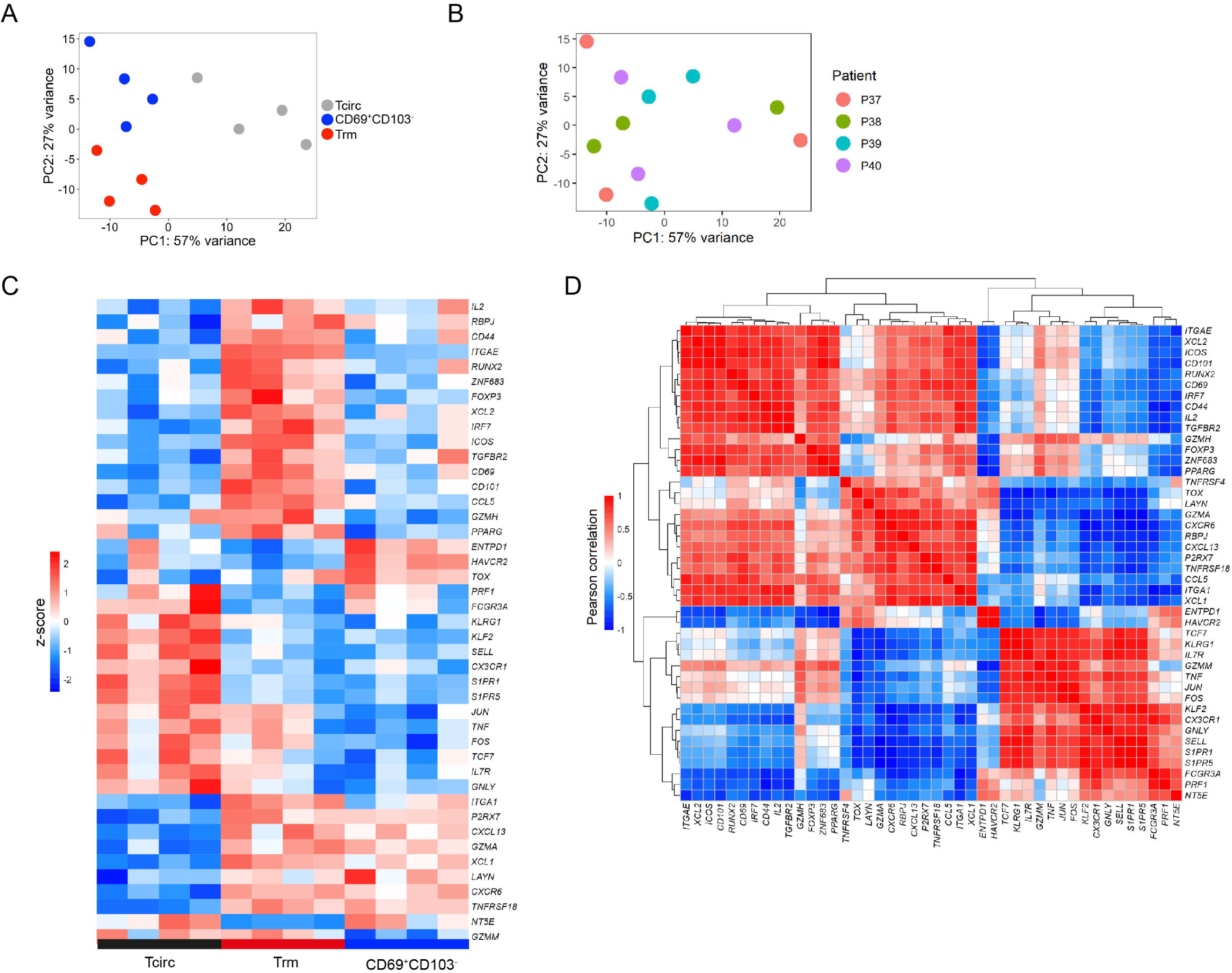
Bulk RNA-seq principal component analysis and transcriptional correlations among memory CD8^+^ T cell subsets. **(A-B)** Principal Component Analysis (PCA) plot showing transcriptomes distribution by memory CD8^+^ TIL subpopulations **(A)** and by patient **(B)**. **(C)** Heatmap showing the expression of key immune genes in each memory CD8^+^ TIL population. **(D)** Pearson coexpression analysis of key immune related genes, values are clustered with complete linkage.

**Supplementary Fig. 4.**
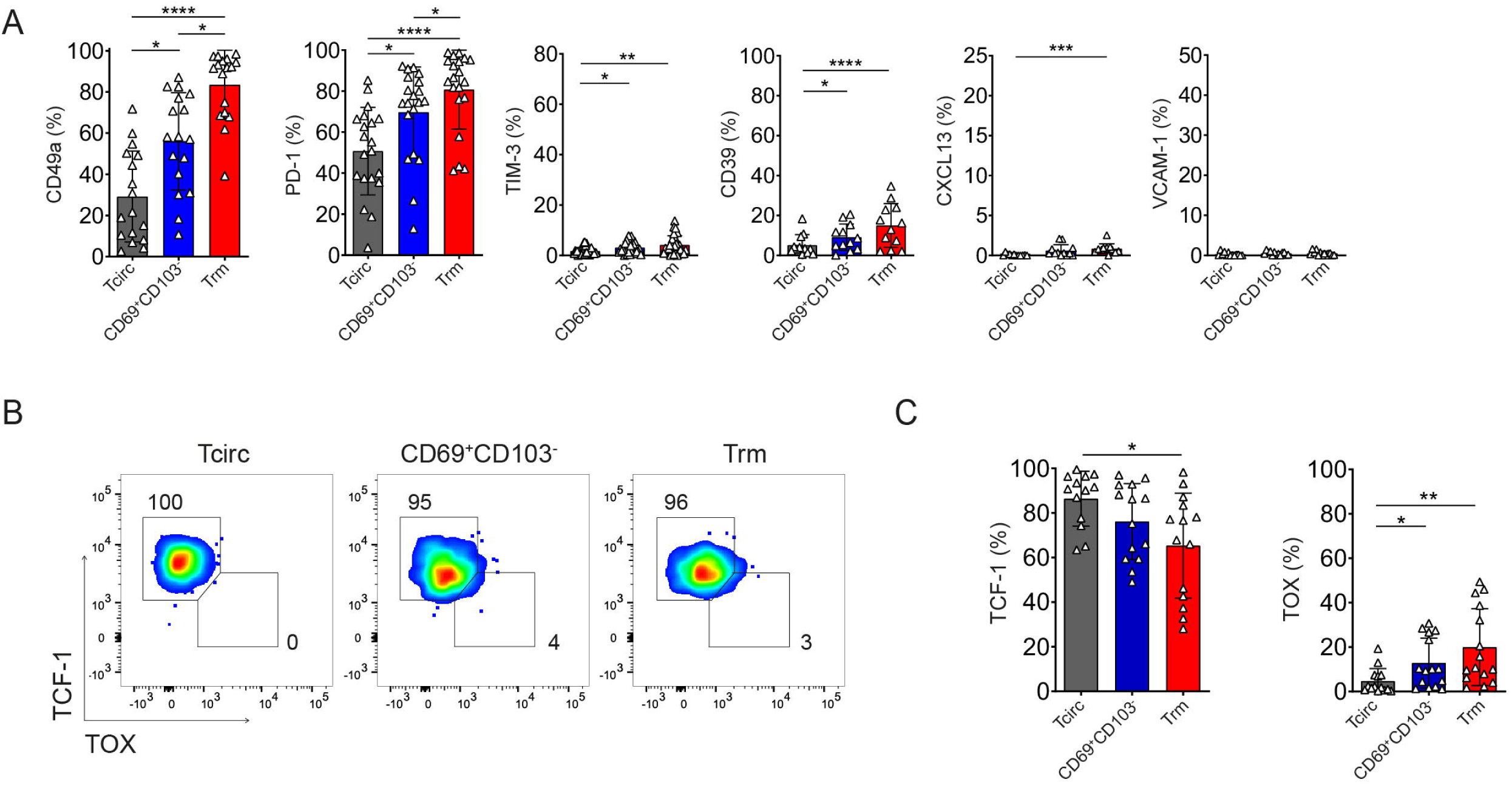
Characterization of non-malignant renal tissue from ccRCC patients by flow cytometry. **(A)** Frequency quantification for the expression of key markers in memory CD8^+^ subpopulations in NMRT. **(B)** FACS plots showing TCF1 and TOX expression measured by intranuclear TF staining and **(C)** frequency quantification of TCF1 and TOX expression in memory CD8^+^ T cells subpopulations from NMRT. Each point represents a patient. Statistical analysis was performed using Friedman paired one-way ANOVA test. *p < 0.05; **p < 0.01; ***p < 0.001; **** p < 0.0001.

**Supplementary Fig. 5.**
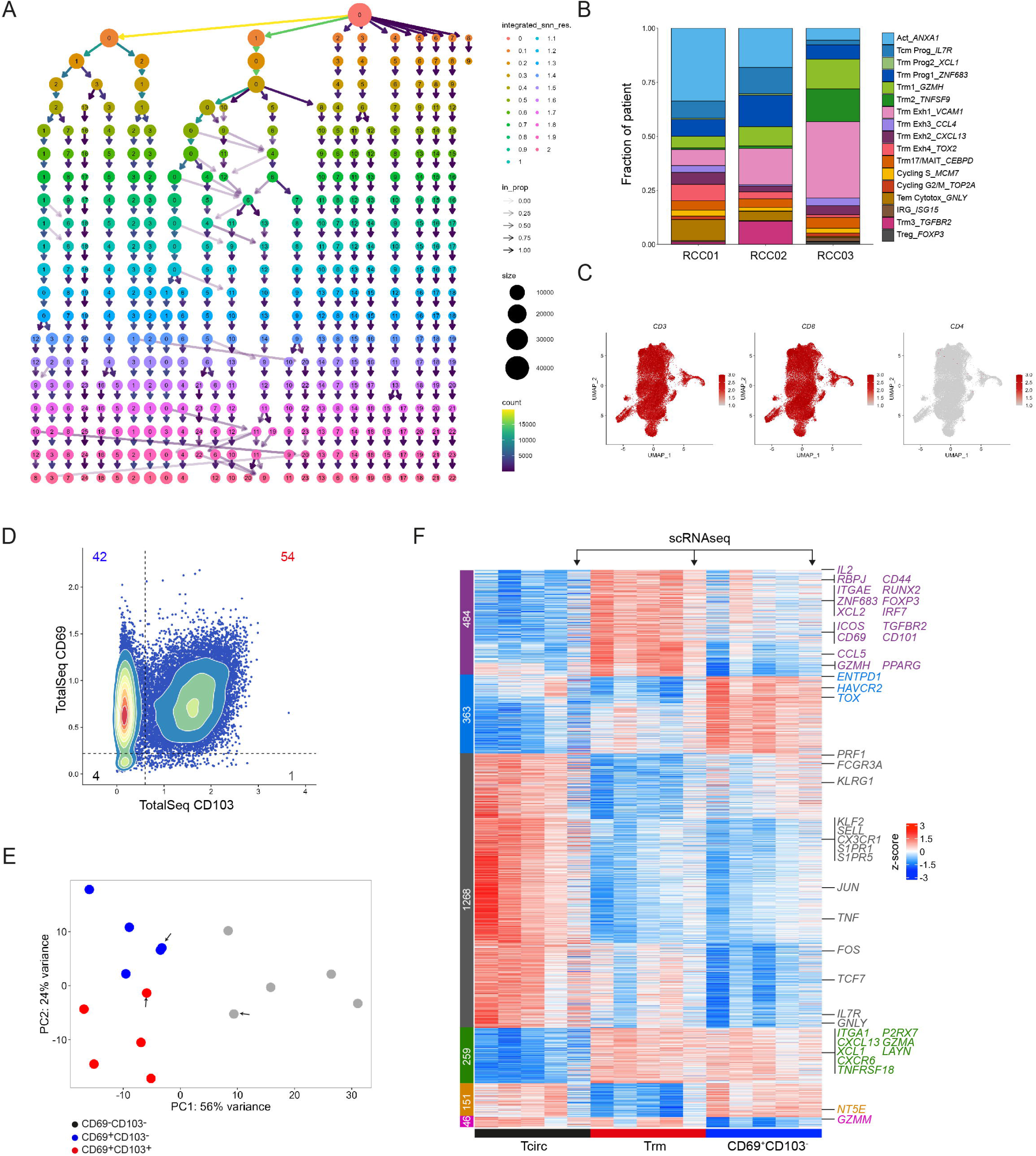
Single-cell quality control, cluster distribution across patients, and ADT-based validation of CD69 and CD103 expression. **(A)** Clustree plot across several scRNA-seq resolutions. **(B)** Bar plot showing cluster distribution in each ccRCC patient. **(C)** UMAP visualization of CD3, CD8 and CD4 normalized gene expression. **(D)** Density plot showing TotalSeq CD69 and CD103 protein expression in memory CD8^+^ T cells from tumor samples. **(E)** Principal Component Analysis (PCA) plot showing transcriptomes (from bulk RNAseq) distribution by memory CD8^+^ TIL subpopulations including pseudo bulk sample generated from scRNAseq data (highlighted with a black arrow). **(F)** Heatmap showing differential expressed genes between Tcirc, CD69^+^CD103^−^ and Trm cells from bulk RNAseq and pseudo bulk sample generated from scRNAseq data. Key genes are labeled. Genes with positive log2-fold change and adjusted p value < 0.05 were considered as upregulated. Genes with negative log2-fold change and adjusted p value < 0.05 were considered as downregulated.

**Supplementary Fig. 6.**
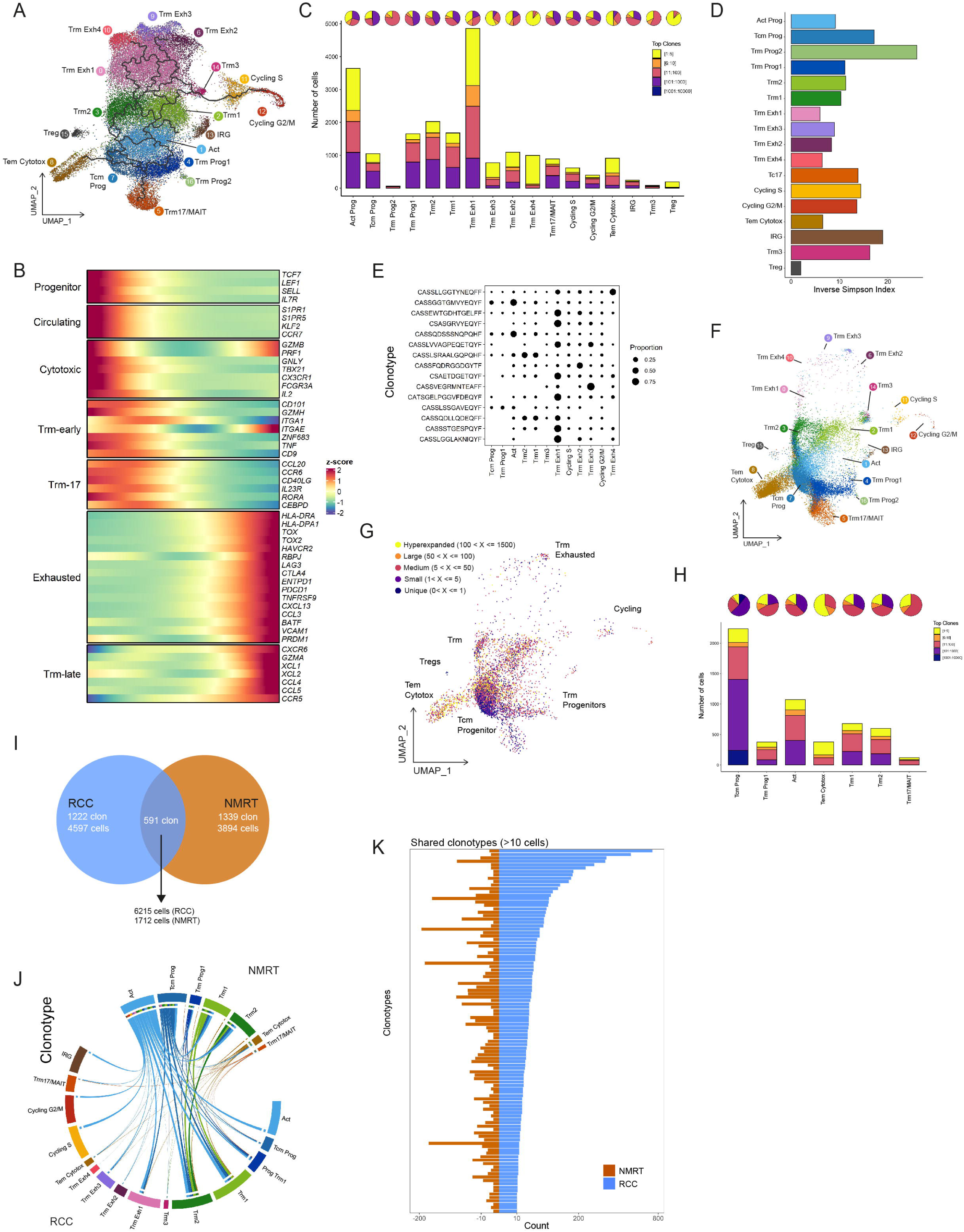
Pseudotime dynamics, gene expression trajectories, and TCR clonal landscape across differentiation states. **(A)** UMAP visualization of the trajectory generated by monocle 3 used to calculate psudotime. **(B)** Heatmap showing key genes normalized expression across the pseudotime. **(C)** Pie charts showing percentages and bar plots showing numbers of cells per scRNAseq clusters from tumor samples colored by Top clones categorization analyzed by scTCRseq **(D)** Barplot showing TCR diversity, measured by inverse Simpson index, for each cluster. **(E)** Bubble plot showing the proportion (dot size) of the top 15 expanded clonotypes in clusters with high Jaccard index. **(F)** UMAP visualization of memory CD8^+^ T cell clusters in the NMRT atlas annotated by label-transfer analysis. UMAP shows 16,615 CD8^+^ T cells from NMRT samples **(G)** UMAP visualization of clonotype frequency categorized as hyperexpanded, large, medium, small, and unique. UMAP shows 5,606 memory CD8^+^ T cells from NMRT cells with TCR data. **(H)** Pie charts showing percentages and bar plots showing numbers of cells per NMRT clusters with more than 100 cells, colored by Top clones categorization. **(I)** Venn diagram showing clonotype (clone) overlap between tumor and NMRT TCR repertoire. **(J)** Circus plot showing clonotype sharing between NMRT clusters and tumor clusters. **(K)** Cell count for clonotypes with more than 10 cells shared between tumor and NMRT.

**Supplementary Fig. 7.**
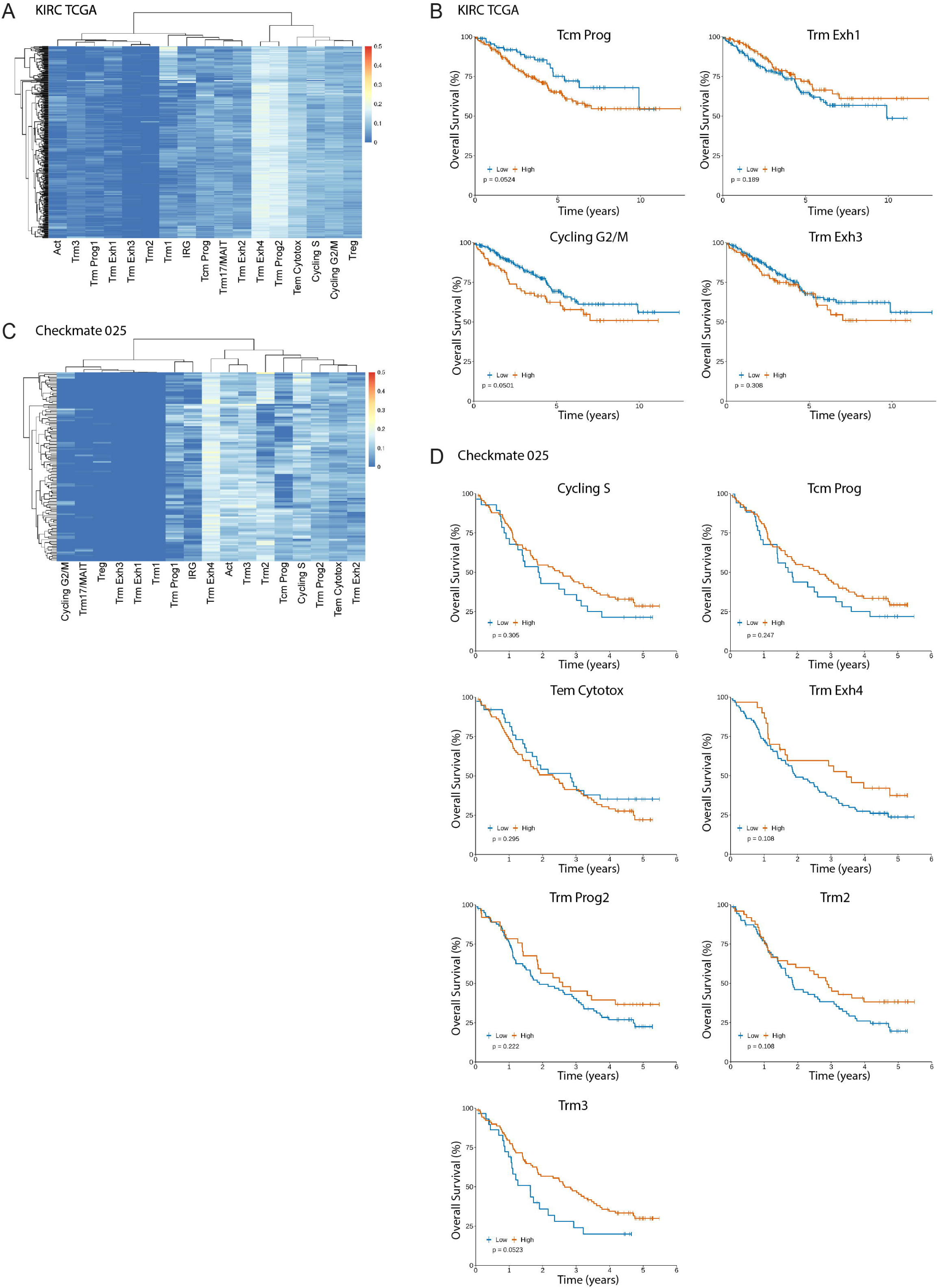
Deconvolution performance and additional survival analyses in TCGA-KIRC and CheckMate-025 cohorts. Single-cell expression matrix was used to deconvolute TCGA-KIRC and Checkmate-025 datasets and survival analysis was performed for each cluster. **(A)** Deconvolution matrix calculated with CIBERSORTx for each cluster present at single-cell analysis for the TCGA-KIRC dataset. **(B)** Kaplan-Meier curves for clusters that showed no significant association with overall survival in the TCGA-KIRC cohort. **(C)** Deconvolution matrix calculated with CIBERSORTx for each cluster present at single-cell analysis for the Checkmate-025 dataset. **(D)** Kaplan-Meier curves for clusters that showed no significant association with overall survival in the CheckMate-025 cohort. In panels **(B)** and **(D)**, patients were stratified into ‘High’ (red) and ‘Low’ (blue) abundance groups for each immune variable using an optimal cut-point, determined by the maximally selected rank statistic. Log-rank test was used to estimate p values.

## References.

1. Hsieh, J. J. et al. Renal cell carcinoma. Nat. Rev. Dis. Primers 3, 17009 (2017).

2. Siegel, R. L., Miller, K. D., Fuchs, H. E. & Jemal, A. Cancer Statistics, 2021. CA Cancer J. Clin. 71, 7–33 (2021).

3. Kim, M.-C., et al. Updates on Immunotherapy and Immune Landscape in Renal Clear Cell Carcinoma. Cancers (Basel). 13, 5856 (2021).

4. Chevrier, S., et al. An Immune Atlas of Clear Cell Renal Cell Carcinoma. Cell 169, 736–749.e18 (2017).

5. Nakano, O., et al. Proliferative activity of intratumoral CD8(+) T-lymphocytes as a prognostic factor in human renal cell carcinoma: clinicopathologic demonstration of antitumor immunity. Cancer Res. 61, 5132–6 (2001).

6. Giraldo, N. A., et al. Orchestration and Prognostic Significance of Immune Checkpoints in the Microenvironment of Primary and Metastatic Renal Cell Cancer. Clinical Cancer Research 21, 3031–3040 (2015).

7. Granier, C., et al. Tim-3 Expression on Tumor-Infiltrating PD-1+CD8+ T Cells Correlates with Poor Clinical Outcome in Renal Cell Carcinoma. Cancer Res. 77, 1075–1082 (2017).

8. Choueiri, T. K., et al. Nivolumab plus Cabozantinib versus Sunitinib for Advanced Renal-Cell Carcinoma. New England Journal of Medicine 384, 829–841 (2021).

9. Motzer, R. J., et al. Nivolumab plus Ipilimumab versus Sunitinib in Advanced Renal-Cell Carcinoma. New England Journal of Medicine 378, 1277–1290 (2018).

10. Motzer, R. J., et al. Avelumab plus axitinib versus sunitinib in advanced renal cell carcinoma: biomarker analysis of the phase 3 JAVELIN Renal 101 trial. Nat. Med. 26, 1733–1741 (2020).

11. Choueiri, T. K., et al. Avelumab + axitinib versus sunitinib as first-line treatment for patients with advanced renal cell carcinoma: final analysis of the phase III JAVELIN Renal 101 trial. Annals of Oncology 36, 387–392 (2025).

12. Zheng, L., et al. Pan-cancer single-cell landscape of tumor-infiltrating T cells. Science (1979). 374, (2021).

13. Chu, Y., et al. Pan-cancer T cell atlas links a cellular stress response state to immunotherapy resistance. Nat. Med. 29, 1550–1562 (2023).

14. Han, J., Khatwani, N., Searles, T. G., Turk, M. J. & Angeles, C. V. Memory CD8+ T cell responses to cancer. Semin. Immunol. 49, 101435 (2020).

15. Luoma, A. M., et al. Tissue-resident memory and circulating T cells are early responders to pre-surgical cancer immunotherapy. Cell 185, 2918–2935.e29 (2022).

16. Ganesan, A.-P., et al. Tissue-resident memory features are linked to the magnitude of cytotoxic T cell responses in human lung cancer. Nat. Immunol. 18, 940–950 (2017).

17. Okła, K., Farber, D. L. & Zou, W. Tissue-resident memory T cells in tumor immunity and immunotherapy. Journal of Experimental Medicine 218, (2021).

18. Krishna, C., et al. Single-cell sequencing links multiregional immune landscapes and tissue-resident T cells in ccRCC to tumor topology and therapy efficacy. Cancer Cell 39, 662–677.e6 (2021).

19. Braun, D. A., et al. Progressive immune dysfunction with advancing disease stage in renal cell carcinoma. Cancer Cell 39, 632–648.e8 (2021).

20. Kourtis, N., et al. A single-cell map of dynamic chromatin landscapes of immune cells in renal cell carcinoma. *Nat*. Cancer 3, 885–898 (2022).

21. Kumar, B. V., et al. Human Tissue-Resident Memory T Cells Are Defined by Core Transcriptional and Functional Signatures in Lymphoid and Mucosal Sites. Cell Rep. 20, 2921–2934 (2017).

22. Yang, K. & Kallies, A. Tissue-specific differentiation of CD8+ resident memory T cells. Trends Immunol. 42, 876–890 (2021).

23. Ye, Q., et al. CD137 Accurately Identifies and Enriches for Naturally Occurring Tumor-Reactive T Cells in Tumor. Clinical Cancer Research 20, 44–55 (2014).

24. Draghi, A., et al. Rapid Identification of the Tumor-Specific Reactive TIL Repertoire via Combined Detection of CD137, TNF, and IFNγ, Following Recognition of Autologous Tumor-Antigens. Front. Immunol. 12, (2021).

25. Carlson, C. M., et al. Kruppel-like factor 2 regulates thymocyte and T-cell migration. Nature 442, 299–302 (2006).

26. Böttcher, J. P., et al. Functional classification of memory CD8+ T cells by CX3CR1 expression. Nat. Commun. 6, 8306 (2015).

27. Mackay, L. K., et al. Hobit and Blimp1 instruct a universal transcriptional program of tissue residency in lymphocytes. Science (1979). 352, 459–463 (2016).

28. Khan, O., et al. TOX transcriptionally and epigenetically programs CD8+ T cell exhaustion. Nature 571, 211–218 (2019).

29. Sekine, T., et al. TOX is expressed by exhausted and polyfunctional human effector memory CD8 ^+^ T cells. Sci. Immunol. 5, (2020).

30. Guo, X., et al. Global characterization of T cells in non-small-cell lung cancer by single-cell sequencing. Nat. Med. 24, 978–985 (2018).

31. Sun, K., et al. scRNA-seq of gastric tumor shows complex intercellular interaction with an alternative T cell exhaustion trajectory. Nat. Commun. 13, 4943 (2022).

32. Savas, P., et al. Single-cell profiling of breast cancer T cells reveals a tissue-resident memory subset associated with improved prognosis. Nat. Med. 24, 986–993 (2018).

33. Gueguen, P., et al. Contribution of resident and circulating precursors to tumor-infiltrating CD8 ^+^ T cell populations in lung cancer. Sci. Immunol. 6, (2021).

34. Li, R., et al. Mapping single-cell transcriptomes in the intra-tumoral and associated territories of kidney cancer. Cancer Cell 40, 1583–1599.e10 (2022).

35. Anadon, C. M., et al. Ovarian cancer immunogenicity is governed by a narrow subset of progenitor tissue-resident memory T cells. Cancer Cell 40, 545–557.e13 (2022).

36. Cook, C. P., et al. A single-cell transcriptional gradient in human cutaneous memory T cells restricts Th17/Tc17 identity. Cell Rep. Med. 3, 100715 (2022).

37. Chung, W., et al. Single-cell RNA-seq enables comprehensive tumour and immune cell profiling in primary breast cancer. Nat. Commun. 8, 15081 (2017).

38. Yang, M., et al. CXCL13 shapes immunoactive tumor microenvironment and enhances the efficacy of PD-1 checkpoint blockade in high-grade serous ovarian cancer. J. Immunother. Cancer 9, e001136 (2021).

39. Liu, B., Zhang, Y., Wang, D., Hu, X. & Zhang, Z. Single-cell meta-analyses reveal responses of tumor-reactive CXCL13+ T cells to immune-checkpoint blockade. *Nat*. Cancer 3, 1123–1136 (2022).

40. Ricketts, C. J., et al. The Cancer Genome Atlas Comprehensive Molecular Characterization of Renal Cell Carcinoma. Cell Rep. 23, 313–326.e5 (2018).

41. Duhen, T., et al. Co-expression of CD39 and CD103 identifies tumor-reactive CD8 T cells in human solid tumors. Nat. Commun. 9, (2018).

42. Giles, J. R., Globig, A.-M., Kaech, S. M. & Wherry, E. J. CD8+ T cells in the cancer-immunity cycle. Immunity 56, 2231–2253 (2023).

43. Russell, J. H. & Ley, T. J. Lymphocyte-Mediated Cytotoxicity. Annu. Rev. Immunol. 20, 323–370 (2002).

44. Niessl, J., et al. Tissue origin and virus specificity shape human CD8 ^+^ T cell cytotoxicity. Sci. Immunol. 10, (2025).

45. Dean, I., et al. Rapid functional impairment of natural killer cells following tumor entry limits anti-tumor immunity. Nat. Commun. 15, 683 (2024).

46. Jonsson, A. H., et al. Granzyme K ^+^ CD8 T cells form a core population in inflamed human tissue. Sci. Transl. Med. 14, (2022).

47. Donado, C. A., et al. Granzyme K activates the entire complement cascade. Nature 641, 211–221 (2025).

48. Schiele, P., et al. CD8 ^+^ T cell–derived CD40L mediates noncanonical cytotoxicity in CD40-expressing cancer cells. Sci. Adv. 11, (2025).

49. Wu, T. D., et al. Peripheral T cell expansion predicts tumour infiltration and clinical response. Nature 579, 274–278 (2020).

50. Corgnac, S., et al. CD103+CD8+ TRM Cells Accumulate in Tumors of Anti-PD-1-Responder Lung Cancer Patients and Are Tumor-Reactive Lymphocytes Enriched with Tc17. Cell Rep. Med. 1, 100127 (2020).

51. Lee, Y. H., et al. IFNγ−IL-17+ CD8 T cells contribute to immunosuppression and tumor progression in human hepatocellular carcinoma. Cancer Lett. 552, 215977 (2023).

52. Lee, M. H., et al. The tumor and plasma cytokine profiles of renal cell carcinoma patients. Sci. Rep. 12, 13416 (2022).

53. Terpstra, M. L., et al. Tissue-resident mucosal-associated invariant T (MAIT) cells in the human kidney represent a functionally distinct subset. Eur. J. Immunol. 50, 1783–1797 (2020).

54. Anvar, M. T., et al. Th17 cell function in cancers: immunosuppressive agents or anti-tumor allies? Cancer Cell Int. 24, 355 (2024).

55. Martin-Orozco, N., et al. T Helper 17 Cells Promote Cytotoxic T Cell Activation in Tumor Immunity. Immunity 31, 787–798 (2009).

56. Chaput, N., et al. Identification of CD8+CD25+Foxp3+ suppressive T cells in colorectal cancer tissue. Gut 58, 520–529 (2009).

57. Kiniwa, Y., et al. CD8+ Foxp3+ Regulatory T Cells Mediate Immunosuppression in Prostate Cancer. Clinical Cancer Research 13, 6947–6958 (2007).

58. Reading, J. L. et al. The function and dysfunction of memory <SCP>CD</SCP> 8 ^+^ T cells in tumor immunity. Immunol. Rev. 283, 194–212 (2018).

59. Koh, C.-H., Lee, S., Kwak, M., Kim, B.-S. & Chung, Y. CD8 T-cell subsets: heterogeneity, functions, and therapeutic potential. Exp. Mol. Med. 55, 2287–2299 (2023).

60. Colak, S. & ten Dijke, P. Targeting TGF-β Signaling in Cancer. Trends Cancer 3, 56–71 (2017).

61. Hao, Y., et al. Integrated analysis of multimodal single-cell data. Cell 184, 3573–3587.e29 (2021).

62. Borcherding, N., Bormann, N. L. & Kraus, G. scRepertoire: An R-based toolkit for single-cell immune receptor analysis. F1000Res. 9, 47 (2020).

63. Newman, A. M., et al. Robust enumeration of cell subsets from tissue expression profiles. Nat. Methods 12, 453–457 (2015).

